# Hypoxia promotes airway differentiation in the human lung epithelium

**DOI:** 10.1101/2024.08.09.607336

**Authors:** Ziqi Dong, Niek Wit, Aastha Agarwal, Dnyanesh Dubal, Jelle van den Ameele, Adam James Reid, James A. Nathan, Emma L. Rawlins

## Abstract

Human early embryos develop under physiological hypoxia, but how hypoxia regulates human organogenesis remains little known. We have investigated oxygen availability effects on the human lung epithelium using organoids. We find first-trimester lung epithelial progenitors remain undifferentiated under normoxia, but spontaneously differentiate towards multiple airway cell types and inhibit alveolar differentiation under hypoxia. Using chemical and genetic tools, we demonstrate that hypoxia-induced airway differentiation is dependent on HIF (Hypoxia-Inducible Factor) pathways, with HIF1α and HIF2α differentially regulating fate decisions. Transcription factors KLF4 and KLF5 are direct targets of the HIF pathway and promote progenitor differentiation to basal and secretory cells. Chronic hypoxia also induces transdifferentiation of human alveolar type 2 cells to airway cells via the HIF pathway, mimicking alveolar bronchiolization processes in lung disease. Our results reveal roles for hypoxia and HIF signalling during human lung development and have implications for aberrant cell fate decisions in chronic lung diseases.

## INTRODUCTION

Human lung development starts around 5 post-conception weeks (pcw) with the emergence of two lung buds from the anterior ventral foregut endoderm.^1,2^ During subsequent branching morphogenesis, the multipotent epithelial progenitors self-renew in the branching tip regions and begin differentiation as they leave the tips.^3–5^ The epithelial tip progenitors initially generate airway (5-17 pcw) and later alveolar (from ∼16 pcw) descendants.^4,6^ However, the maternal-placental blood circulation remains incomplete until the end of the first trimester (∼12 pcw), when the endovascular extravillous trophoblasts remodel the spiral arteries to release blood into the extravillous space.^7^ Consequently, the oxygen tension within first-trimester placenta can be as low as 10-40 mmHg (approximately 1.3-5.3% of standard atmospheric pressure).^8^ This results in a highly hypoxic environment during early human organogenesis, including initial airway formation stage. However, any direct effects of this physiological hypoxia on human lung epithelial development remain unknown.

The Hypoxia-Inducible Factor (HIF) pathway orchestrates diverse cellular responses to hypoxia, including cell proliferation, differentiation, and metabolic reprogramming.^9–12^ The activity of HIF pathways is primarily regulated by post-translational control of HIFα (HIF1α; HIF2α, also known as EPAS1) subunits.^13^ Briefly, in normoxia, HIFα is hydroxylated by prolyl hydroxylase enzymes (PHDs), then ubiquitinated by the Von Hippel–Lindau (VHL) E3 ligase followed by proteasome degradation.^14–16^ When oxygen is limited, intact HIFα translocates into the nucleus and forms heterodimers with HIF1β (also known as ARNT). The HIF complexes bind Hypoxia-Response Elements (HREs) and activate downstream genes with various co-activators/chromatin modifiers.^13^

Hypoxia and the HIF pathway can modulate the processes of lung development, repair and disease.^17,18^ Hypoxia induced *Drosophila* larva tracheal sprouting through Sima (HIFα homolog in *Drosophila*).^19–21^ Mild hypoxia increased branching of mouse embryonic lung explants.^22,23^ Hypoxia can also induce neuroendocrine (NE), or goblet cell differentiation in the mouse or human adult airway epithelium respectively.^24,25^ However, the functions and direct targets of HIFα homologs governing various lung epithelial lineages remain largely unknown. In the alveoli, alveolar type 2 (AT2) cells are adult stem cells responsible for restoring alveolar epithelium after mild injuries.^26–28^ After severe damage, however, ectopic basal cells can emerge in the alveolar epithelium impairing gas exchange capacity.^29–36^ In mice this is likely via HIF1α activation of *Krt5* transcription in lineage-negative progenitors.^28^ In human fibrotic lungs, aberrant basal-like and other airway cells accumulate in distal lungs and exhibit hypoxia signatures.^32–36^ Improved understanding of cellular responses to hypoxia, particularly the functions and mechanisms of the HIF pathway in human lung epithelial cell fate decisions, will facilitate the development of intervention methods for hypoxia-induced lung damage or aberrant remodelling.

Here we have used human fetal lung-derived organoid models to elucidate the direct effects of hypoxia on human lung epithelial progenitors. First-trimester lung progenitors cultured in a self-renewing condition spontaneously differentiated to airway fate under hypoxia, and simultaneously repressed alveolar-specific genes. Activation of the HIF pathway recapitulated these phenotypes. We systematically dissected the functions of HIF1α and HIF2α and identified KLF4 and KLF5 as direct downstream targets, mediating progenitor differentiation to basal and secretory cells. Hypoxia also reprogrammed differentiated human AT2 cells to airway cells in a HIF-dependent manner, suggesting major conservation of HIF functions between the embryonic progenitors and adult AT2 cells. Therefore, hypoxia emerges as a developmental cue directly promoting airway differentiation of fetal lung epithelial progenitors, with implications for understanding aberrant transdifferentiation of AT2 cells in chronic lung disease.

## RESULTS

### Hypoxia induces airway differentiation of human fetal lung epithelial progenitors grown in self-renewing conditions

Human fetal lung multipotent tip epithelial progenitors can be isolated and expanded *in vitro*.^4,37^ To study the direct effects of hypoxia on these epithelial progenitors, we first cultured dissected fetal lung tips (7-9 pcw) as organoids in self-renewal medium (SRM). After expanding for 1-2 passages, we dissociated the organoids to single cells and divided the progenitor pool into either normoxia (20-21% O_2_, 5% CO_2_) or hypoxia (2% O_2_, 5% CO_2_) conditions (Figure 1A). To evaluate cell identity, we monitored organoid morphology (Figures 1B and 1C) and gene expression of progenitor and differentiation markers via RT-qPCR (Figure 1D). Organoids cultured under normoxia required frequent passaging (every 6-7 days) and sustained largely unchanged expression of key marker genes across multiple passages (Figure S1A). Whereas progenitors cultured under hypoxia (2% O_2_) slowed their proliferation (passaging every 2 weeks) and acquired folded morphologies (Figures 1B and 1C). Moreover, hypoxic culture for 15 or 30 days consistently elevated the expression levels of airway cell markers despite organoid background variations, including for basal cells (*TP63, KRT5*), secretory cells (*SCGB3A2*) and neuroendocrine cells (*ASCL1, GRP*), and concurrently downregulated the canonical AT2 marker (*SFTPC*) (Figure 1D). The emergence of airway cell markers was confirmed by immunostaining (Figure 1E). Although hypoxia slightly increased *SOX2* and *SOX9* at the mRNA level (Figure 1D), SOX2 protein (expressed in early human lung tip, stalk, and airway cells) remained, while most cells lost SOX9 (tip cell specific) under hypoxia (Figure 1E), consistent with SOX2 functioning in airway differentiation.^4,37–39^ Extended hypoxia culture (2% O_2_ for 45 days) decreased expression of the proliferation marker (Ki67) without altering lung cell identity (NKX2.1) (Figure 1E).

**Figure 1.**
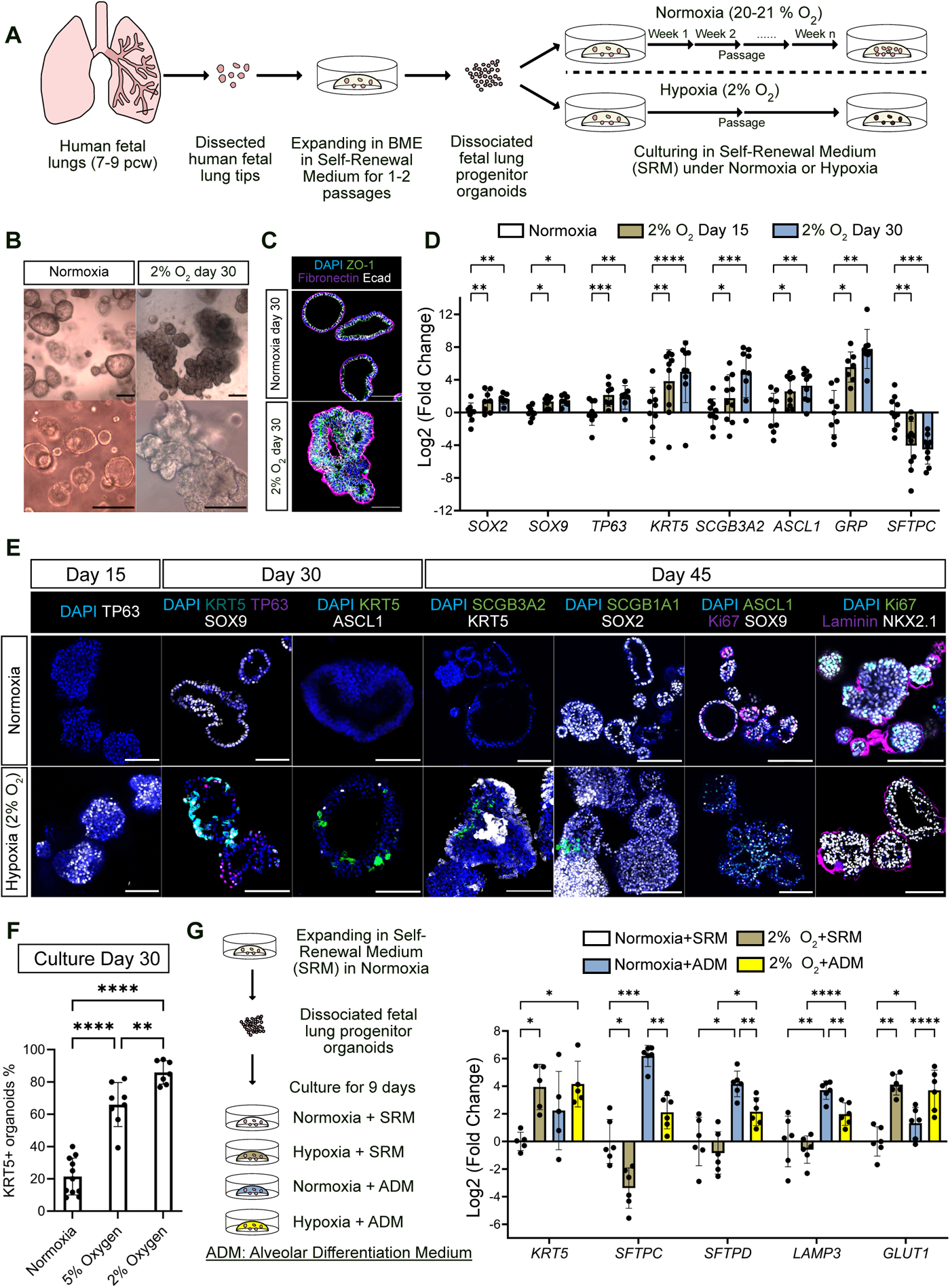
Hypoxia promotes airway differentiation of first-trimester human fetal lung epithelial progenitors. (A) Experiment design. The epithelial progenitors were isolated from human fetal (7-9 pcw) lung tips, expanded in SRM, and treated with normoxia or hypoxia (2% O_2_). (B) Brightfield images of lung progenitor organoids cultured under normoxia or hypoxia for 30 days. (C) Hypoxia induced organoid shape changes visualised by ZO-1 (Zonula Occludens-1), Fibronectin and Ecad (E-cadherin). (D) Progenitor and differentiation marker gene expression under normoxia, or hypoxia (2% O_2_) for 15 and 30 days detected by RT-qPCR. Fold changes were normalised to the mean of the normoxia samples. Bars represent mean Log_2_(fold change) ± standard deviation (SD), n = 10 experimental replicates from 8 biological donors. Statistical test: two-way ANOVA with Geisser-Greenhouse correction and Dunnett’s multiple comparisons test. (E) Immunostaining for progenitor (SOX9, SOX2), basal cell (TP63, KRT5), secretory cell (SCGB3A2, SCGB1A1), neuroendocrine cell (ASCL1), proliferation (Ki67), lung identity (NKX2.1) markers and extracellular matrix protein (Laminin) in organoids cultured under normoxia, or hypoxia (2% O_2_) for 15, 30 and 45 days. DAPI: nuclei. Representative images from 4 organoid lines. (F) The percentage of organoids containing ≥ 1 KRT5^+^ cell(s). The average values were calculated from multiple independent experiments n = 11 (normoxia), 8 (5% O_2_), 7 (2% O_2_) from 3 biological donors. Data shown as mean ± SD. Statistical test: one-way ANOVA with Tukey’s multiple comparisons test. (G) RT-qPCR of organoids cultured in normoxia + SRM, normoxia + ADM, hypoxia + SRM, and hypoxia + ADM for 9 days. The fold changes were normalised to the mean of normoxia + SRM condition. Bars represent mean Log_2_(fold change) ± SD, n = 6 biological donors. Statistical test: two-way ANOVA with Tukey’s multiple comparisons test. Scale bars = 100 µm all panels. Gene expression was normalised to *ACTB* in RT-qPCR. Significance levels: *p < 0.05, **p < 0.01, ***p < 0.001, ****p < 0.0001. See also Figure S1.

To examine the dependence of the phenotype on hypoxia levels, we exposed progenitor organoids to 5% O_2_. At 5% O_2_ the organoids also upregulated airway and downregulated AT2 cell markers, but to a lesser extent than at 2% O_2_ (Figure S1B). To further compare the effects of hypoxia levels, we cultured organoids from the same dissociated progenitor pool under normoxia, 2% O_2_ or 5% O_2_ for 30 days and detected KRT5^+^ basal cells by immunostaining. Both 2% O_2_ and 5% O_2_ conditions promoted KRT5^+^ cells (Figures 1E and S1C). However, the percentage of organoids containing ≥ 1 KRT5^+^ cells was highest under 2% O_2_ (Figure 1F). We selected 2% O_2_ in subsequent experiments to mimic the hypoxia condition during first-trimester lung development and to enhance the hypoxia-induced differentiation phenotype.

To test the hypoxia effects on progenitor differentiation towards alveolar lineages, we cultured cells in either SRM or a published alveolar differentiation medium (ADM)^40^ under normoxia or hypoxia (2% O_2_) for 9 days. In the SRM condition, 9-day hypoxia was sufficient to induce *KRT5* and the hypoxia-responsive gene *GLUT1* (also known as *SLC2A1*), while decreasing *SFTPC* (Figure 1G). The ADM condition promoted AT2 markers (*SFTPC, SFTPD, LAMP3*) under normoxia. However, hypoxia diminished ADM’s effects by inhibiting AT2 marker gene expression and promoting *KRT5* (Figure 1G). To test if the hypoxia-induced differentiation effects are evolutionarily conserved, we isolated mouse lung tip progenitors during the branching stage (between E11.5-E14.5) and cultured them in a published self-renewal medium.^41^ Hypoxia promoted mouse airway marker gene expression (*Foxj1, Sox2*) but did not significantly decrease an AT2 marker (*Sftpc*) (Figure S1D). Collectively, these data illustrate that hypoxia promotes airway fate while limiting alveolar fate in human lung epithelial progenitors.

### Concurrent emergence of basal, neuroendocrine and secretory cell lineages under hypoxia

To determine the cellular dynamics underlying hypoxia-induced progenitor differentiation, we conducted a time-series single-cell RNA-sequencing experiment. We sampled organoids cultured under normoxia and 8, 16, 24, and 32 days of hypoxia from two fetal lungs (9 pcw) and processed together to minimise batch effects (Figure 2A). Combining all samples yielded a 65,475-cell transcriptomic dataset with > 4,200 median number of genes per cell (Figure S2A). Overall, we identified 11 cell populations: three populations of progenitors (designated as tip progenitors, primed progenitors and airway progenitors), differentiated airway cell types (basal cells, neuroendocrine cells, secretory-like cells, and hillock-like cells), cycling cells and two intermediate populations (Figure 2B). The two donor replicates were highly consistent as visualised by Uniform Manifold Approximation and Projection (UMAP) (Figure S2B) and had similar contributions to most cell types (Figure S2C). We therefore merged the data from both organoid lines for analysis.

**Figure 2.**
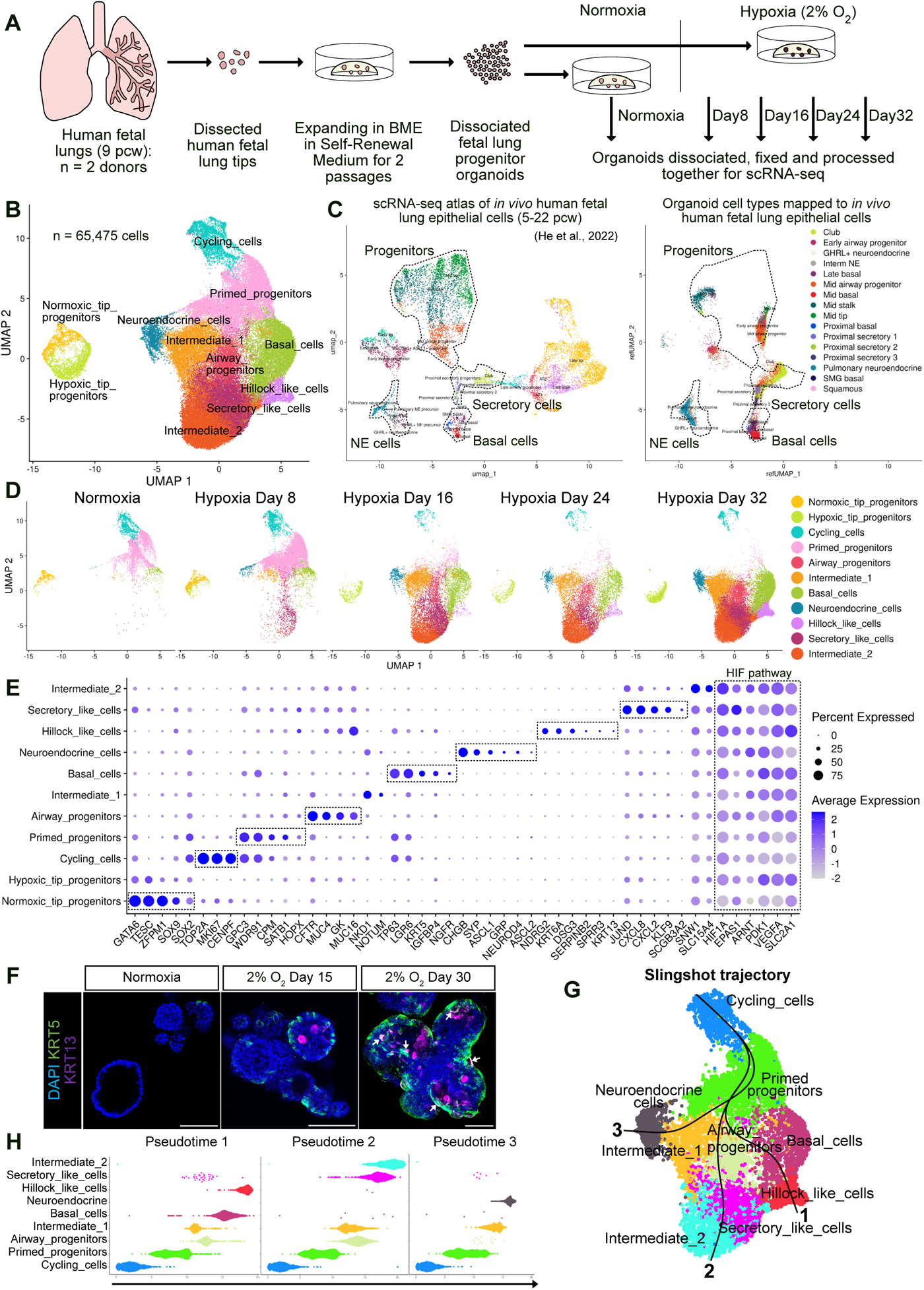
Emergence of basal, neuroendocrine, secretory-like and hillock-like cells under hypoxia. (A) Experiment design. The epithelial progenitor organoids were derived from two human fetal (9 pcw) lungs and treated with normoxia or hypoxia (2% O_2_) in SRM. The organoids were sampled, dissociated, and fixed over several days, and processed together for library preparation. (B) UMAP of cells from all samples with identified cell types. (C) Left panel: reference UMAP of human fetal lung epithelial cell atlas.^6^ Right panel: all cells from the organoids projected to the reference UMAP. (D) The organoid cells sampled at different days shown in the UMAP in (B) with annotated cell types. (E) Expression patterns of canonical *in vivo* lineage markers, cell type-specific markers newly identified from the organoid dataset, and HIF pathway-related genes. (F) Hillock-like cells (KRT13^+^) emerged at day 15 and day 30 in hypoxic organoids. KRT5: basal cell marker. Arrows indicate KRT5^+^KRT13^+^ cells. Scale bars = 100 µm. (G) and (H) Slingshot trajectory analysis. (G) Three pseudotime trajectories started from cycling cells and diverging at primed progenitors. (H) Cell type changes along the trajectories. See also Figures S2 and S3.

We benchmarked the organoid data against *in vivo* human fetal lung epithelial cells by projecting organoid cells onto a reference atlas UMAP.^42,6^ Consistent with the RT-qPCR data (Figure 1D), organoid cells mainly mapped to mid-stage tip/stalk cells, airway progenitors and differentiated airway cells (basal cells, neuroendocrine cells and secretory cells; note that the hillock cells were not captured in the reference atlas), but not to later developmental stage tip progenitors or alveolar cells (Figure 2C). The gene capture rate was greater than we have previously achieved with scRNA-seq,^6^ revealing unexpected heterogeneity in the normoxic self-renewing organoids (Figure 2D, left panel). Normoxic organoids consisted of two progenitor populations, cycling cells, and small numbers of differentiating cells (Figures 2D, S2D and S2E). We designated these progenitors as ‘tip’ and ‘primed’. Tip progenitors were *SOX9*^high^*SOX2*^low^ and expressed human lung tip-enriched markers *GATA6* and *TESC* (Figures 2E and S2F), with high regulon activity for SOX9 and GATA6 (Figure S3A).^4,6,37,39,43,44^ Primed progenitors were *SOX9*^low^*SOX2*^high^ with a subset expressing low levels of *HOPX*, a lung stalk cell feature, and *TP63*, a basal cell transcription factor (Figures 2E and S2F).^6^ Primed progenitors expressed both epithelial progenitor marker (*CPM*) and developmental genes (*WDR91*, *SATB1*) with high FOXA1 regulon activity, indicating a differentiation-priming state (Figures 2E and S3A).^45^ The tip and primed progenitors in organoids likely represent tip cells and stalk (or tip-adjacent) cells *in vivo*. These data are consistent with our previous lower-resolution scRNA-seq data of tip-derived organoids in which the cells mapped to the tip/stalk boundary of the human fetal lung.^6^

During hypoxia culture, proliferating cell numbers decreased, and organoid cell composition changed, particularly between days 8 and 16, with normoxic tip and primed progenitors nearly disappearing (Figures 2D and S2E). Most cell populations that predominated in hypoxia could be assigned to known *in vivo* cell types. Airway progenitors highly expressed *CFTR*, the *in vivo* airway epithelial progenitor marker,^46–48^ as well as *MUC4* and *MUC16*, the proximal secretory cell markers.^6^ Basal cells (*TP63*, *KRT5*, *LGR6*, *NGFR*) and neuroendocrine cells (*ASCL1*, *GRP*, *CHGB*, *SYP*, *ASCL2*, *NEUROD4*) expressed canonical markers (Figures 2D, 2E, and S2F). Surprisingly, we identified a hillock-like cell population, marked by *KRT6A*, *KRT13*, *DSG3*, *SERPINB2*, and *SPRR3*, that appeared from hypoxia day 16 (Figures 2D, 2E, S2E, and S2F).^49–52^ We confirmed the existence of hillock-like cells by detecting KRT13^+^ cells specifically in the hypoxic organoids (Figure 2F). The secretory-like cells expressed *SCGB3A2* and chemokine genes *CXCL8* and *CXCL2* which were enriched in proximal secretory cells *in vivo*.^6^ The secretory-like cells also highly expressed the oxidative stress-responsive gene *JUND* with activated c-Jun N-terminal kinase (JNK) and NF-κB pathways, showing a cell type-specific response to hypoxia (Figures 2E, S2F and S3A).^53,54^ The cycling cells had mixed identities and persisted under hypoxia, consistent with the slow expansion phenotype of hypoxic organoids (Figures 2E, S2E and S2F).

To infer relationships between different cell populations, we conducted trajectory analysis with Slingshot.^55^ By defining the starting point of the pseudotime as cycling cells, we found that the trajectory branched at primed progenitors (Figure 2G). One branch led to basal cells and hillock-like cells, and the second branch led to airway progenitors, secretory-like cells and intermediate-2 cells. The third branch led to neuroendocrine cells through intermediate-1 cells (Figures 2G and 2H). This early trajectory branching event at the primed progenitors corresponded well with the cell population emergence order along actual sampling time points (Figure 2D), and a similar analysis performed with a complementary method, Monocle 3 (Figures S3B and S3C).^56^ Consistent with the primed progenitors being the source of the differentiated cells, this population is enriched in developmental genes associated with morphogenesis (Figures S3D and S3E). Interestingly, the tip and primed progenitor populations responded to hypoxia differently. In contrast to the hypoxia-induced airway differentiation of the primed progenitors, in hypoxia the tip progenitors downregulated genes related to the cell cycle and upregulated genes involved in cell adhesion (Figures S3F and S3G), but did not express differentiation markers (Figures 2E and S2F).

Canonical HIF-pathway genes (*PDK1*, *VEGFA/VEGF*, *SLC2A1/GLUT1*) were highly expressed in hypoxic tip progenitors, airway progenitors and other intermediate and differentiated cells that emerged under hypoxia (Figure 2E).^13^ Moreover, the HIF genes themselves were differentially expressed within our scRNA-seq data. *HIF1A* was more broadly expressed than *HIF2A* (*EPAS1*) and *ARNT* (*HIF1B*), though they all had higher expression levels in neuroendocrine cells, hillock-like cells, secretory-like and intermediate 2 cells. *HIF1A* also showed higher expression levels than *HIF2A* in primed progenitors and basal cells (Figures 2E and S2F). The differential expression levels of HIF-pathway genes suggest cell type-specific roles of HIF1α and HIF2α.

### The HIF pathway is activated in hypoxia and is sufficient to drive progenitor differentiation

To analyse the HIF pathway activity in progenitor organoids, we used the HRE-ODD-GFP reporter construct.^57^ The Oxygen-Dependent Degradation (ODD) domain-tagged GFP is expressed when stabilised HIF-1/2 complexes bind to the HRE sequence under hypoxia, and the GFP protein is subsequently degraded if the oxygen level increases (Figure 3A). Under normoxia, frequent organoid passaging (routine laboratory practice) maintains the HIF-responsive GFP^+^ cells at a baseline level (Figures 3B and 3C). However, GFP^+^ organoids (organoids containing ≥ 1 GFP^+^ cell) accumulated significantly when organoids were grown in normoxia without timely passaging, likely due to local hypoxia caused by crowding and increased oxygen consumption. In contrast, GFP was rapidly activated upon hypoxia exposure. In hypoxia, the GFP^+^ organoid proportion peaked at day 4 then gradually decreased (Figures 3B and 3C), indicating temporal regulation of the HIF pathway. However, Stabilised HIF1α and HIF2α proteins were detected throughout the 1-month hypoxia culture (Figure S4A), suggesting additional mechanisms regulating HIF transactivation activity. Moreover, consistent with rapid HIF pathway activation, the expression of canonical HIF target genes *VEGF* and *GLUT1*, as well as *HIF2A* and *KRT5* increased within 1-week hypoxia culture (Figure S4B).

**Figure 3.**
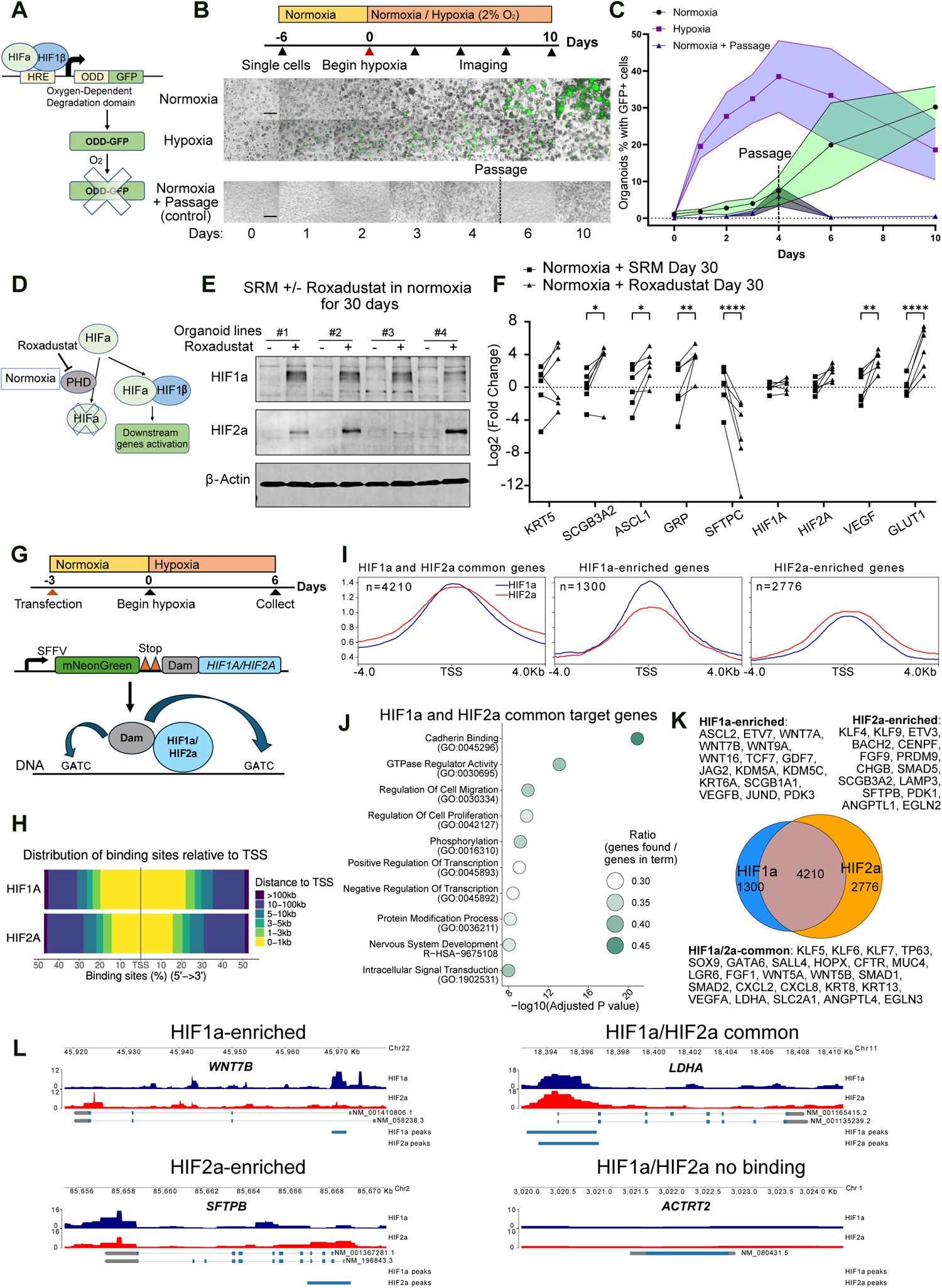
The HIF pathway is activated in hypoxic lung organoids. (A) Diagram of HRE-ODD-GFP reporter construct and oxygen-dependent GFP turnover. (B) Microscope images of merged brightfield and GFP channels. The HRE-ODD-GFP reporter progenitors were cultured under normoxia for 6 days to form organoids, then treated with normoxia or hypoxia for 10 days without passaging. Control cells were cultured under normoxia with routine passaging. Scale bars = 600 µm. (C) The percentage of organoids containing ≥ 1 GFP^+^ cell(s). Data shown as mean ± SD, n = 7 (normoxia), 8 (hypoxia) experimental replicates from 2 biological donors. (D) Roxadustat (FG-4592) inhibits PHD enzymes under normoxia and stabilises HIFα subunits. (E) HIF1α and HIF2α were stabilised under normoxia by Roxadustat in 4 organoid lines with β-actin as loading control. (F) Roxadustat treatment activated the HIF pathway under normoxia and recapitulated hypoxia-induced differentiation. RT-qPCR detection of organoids cultured in SRM ± Roxadustat for 30 days. Fold changes were normalised to the mean of SRM – Roxadustat (with DMSO) condition. Data shown as Log_2_(fold change), n = 6 biological donors. Statistical comparisons by two-way ANOVA with Bonferroni’s multiple comparisons test. (G) Design of targeted DamID-seq for HIF1α and HIF2α. Dam-HIFα fusion proteins are expressed at a low level due to rare translation reinitiation events. The fusion proteins methylate adenines in the GATC sequences near their DNA binding sites. (H) Global distributions of HIF1α and HIF2α binding sites relative to the Transcriptional Start Site (TSS). (I) Quantification of HIF1α and HIF2α binding signals surrounding the TSS. HIF1α and HIF2α signals were normalised to Dam-only control. (J) Gene ontology analysis of HIF1a and HIF2a common target genes. (K) Venn diagram comparing HIF1α and HIF2α target genes with highlighted gene lists. (L) Gene track views showing averaged DamID signals and consensus peaks from three biological replicates over selected HIF1α and HIF2α binding and no binding genes. Gene expression was normalised to *ACTB* in RT-qPCR. Significance levels: *p < 0.05, **p < 0.01, ***p < 0.001, ****p < 0.0001. See also Figure S4.

To test whether activation of the HIF pathway alone was sufficient to drive airway differentiation, we applied Roxadustat (FG-4592), a PHD inhibitor, to progenitor organoids under normoxia (Figure 3D). Both HIF1α and HIF2α were stabilised in normoxia by Roxadustat treatment (Figure 3E). Moreover, Roxadustat treatment in SRM for 30 days under normoxia increased airway cell markers (*KRT5*, *SCGB3A2*, *ASCL1*, *GRP*) and HIF target genes (*VEGF*, *GLUT1*), but decreased the AT2 marker *SFTPC* (Figure 3F), recapitulating the hypoxia-induced differentiation phenotype. These data strongly suggest that the hypoxia-induced airway differentiation is mediated by the HIF pathway.

As hypoxia or Roxadustat treatment stabilised both HIF1α and HIF2α proteins, preventing from distinguishing HIF-1 and HIF-2 effects, we used targeted DamID (DNA adenine methyltransferase identification)-sequencing to compare the genomic binding sites of HIF1α and HIF2α. Targeted DamID has been employed to map SOX9 and NKX2.1 binding sites in lung organoids.^40,43^ Here, the lung progenitors derived from three donors were transduced by Dam-only control, Dam-HIF1α or Dam-HIF2α fusion proteins and treated with hypoxia for 6 days (Figure 3G). From principal component analysis (PCA), the Dam control samples clustered together and separated from Dam-HIFα samples, while Dam-HIF1α and Dam-HIF2α partially overlapped (Figure S4C). We normalised Dam-HIF1α and Dam-HIF2α signals to Dam control across the genome to calculate enrichment levels of HIF1α and HIF2α binding. We defined the peaks as consensus peaks only if they existed in all three replicates. HIF1α and HIF2α consensus peaks were mostly enriched in TSS (transcription start site) or promoter-adjacent regions (Figures 3H and S4D). By assigning consensus peaks to the nearest TSS, we identified HIF1α and HIF2α target genes. Among them, HIF1α and HIF2α shared 4210 target genes that were involved in cell migration, proliferation, transcription and protein modification processes (Figures 3I-K). There were also HIF1α-enriched (1300 genes) and HIF2α-enriched (2776 genes) target genes, that were preferentially bound by either factor (Figure 3I). These included different lineage markers and signalling pathways (Figures 3K and 3L). Dam-HIF1α and Dam-HIF2α had enriched signals at canonical hypoxia-responsive genes (*LDHA*, *ANGPTL4*, *VEGFA*) (Figures 3L and S4E). HIF1α and HIF2α also differentially bound lineage markers (*SFTPB*, *MUC4*, *HOPX*, *KRT6A*, *CXCL2*, *ASCL2*, *LGR6*) and development-related transcription factors (*SALL4*, *KLF7*) (Figures 3L and S4E), overlapping with cell type-specific markers and regulons in the organoid scRNA-seq dataset (Figures 2E, S2F, and S3A). These results suggested HIF1α and HIF2α had both common and distinct functions in hypoxic lung organoids.

### HIF1α is required for hypoxia-induced airway differentiation

We used an inducible CRISPRi system to interrogate HIF1α and HIF2α functions in progenitor fate decisions separately.^58^ With previously evaluated gRNAs targeting *HIF1A*,^59^ the CRISPRi system efficiently knocked down HIF1α in hypoxia within 4 days (Figure 4A). To examine how depleting HIF1α affects progenitor differentiation, we cultured non-targeting control (NTC) or *HIF1A*-targeting gRNA transfected organoids under hypoxia (2% O_2_) for 30 days. Depletion of *HIF1A* limited the hypoxia-induced expression of airway cell markers (*KRT5*, *SCGB1A1*, *ASCL1*, *GRP*) (Figure 4B). This was confirmed for KRT5 and ASCL1 at the protein level (Figure 4C). Intriguingly, HIF1α-knockdown consistently resulted in even lower *SFTPC* levels than the non-targeting control (Figure 4B).

**Figure 4.**
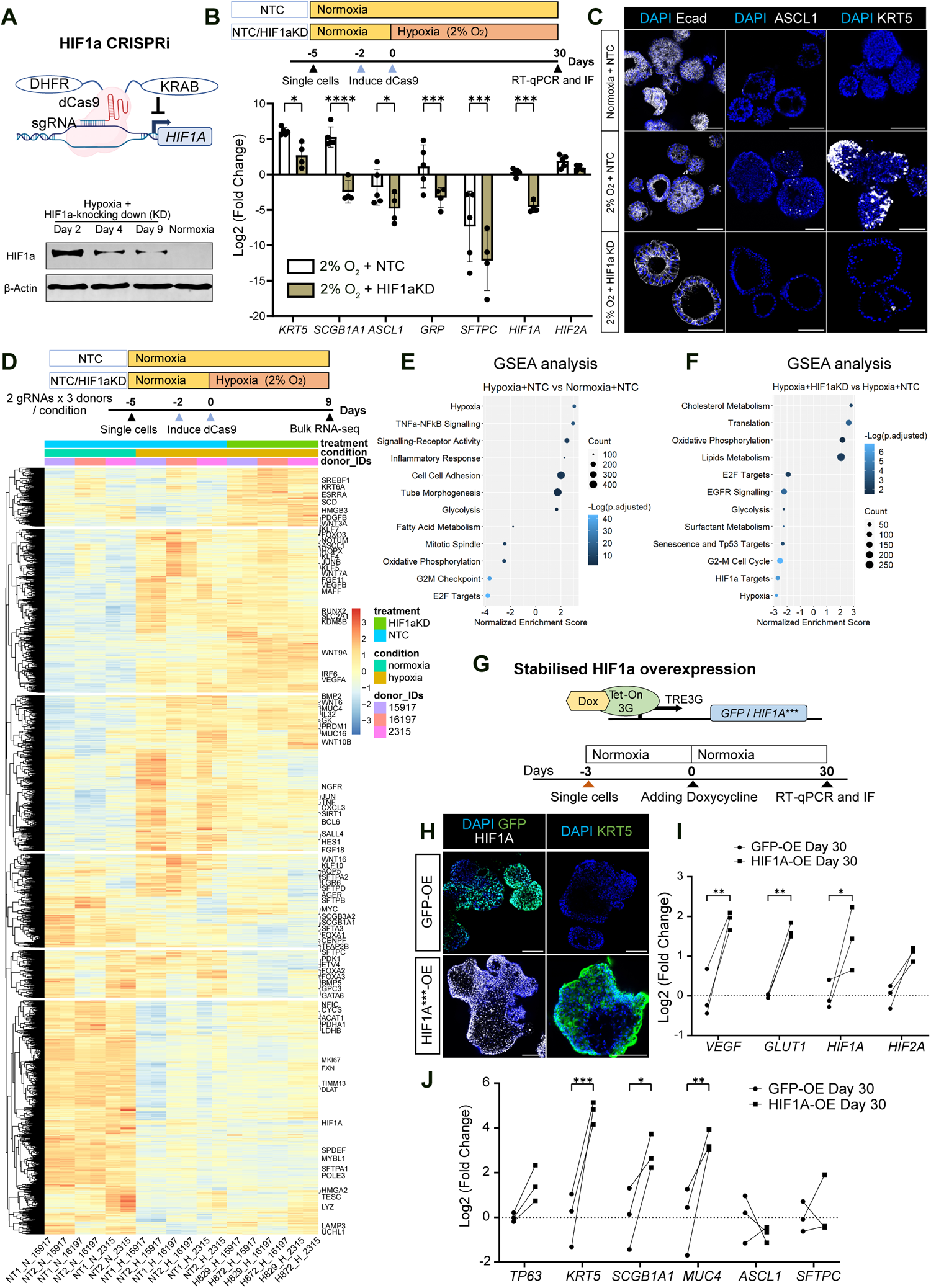
HIF1α is required for hypoxia-induced airway differentiation. (A) HIF1α is inhibited by CRISPRi. The dCas9-KRAB effector was tagged with DHFR and stabilised in presence of TMP (Trimethoprim) to reduce leaky expression from the Tet-on promoter. HIF1α protein levels decreased after 4-9 day knock down under hypoxia as shown by Western blot. (B) Upper panel: experimental design. The non-targeting control (NTC) or *HIF1A-*knock down (*HIF1A-* KD) organoids were cultured under normoxia or hypoxia for 30 days. The dCas9 was induced 2 days ahead of hypoxia treatment. Lower panel: RT-qPCR results. Fold changes were normalised to the average of NTC + normoxia condition (not shown). Bars represent mean Log_2_(fold change) ± SD, n = 4 experimental replicates from 3 biological donors. 2 gRNAs tested. (C) Immunostaining of organoids with NTC or *HIF1A-*KD induction for 30 days showed changes in organoid shape (Ecad: E-cadherin) and differentiation (ASCL1, KRT5). Representative images from 3 organoid lines. (D) The NTC and *HIF1A*-KD organoids were cultured under normoxia or hypoxia for 9 days and used for bulk RNA-seq with 2 gRNAs and 3 biological donors for each condition. Heatmap showing 7309 differentially expressed genes (DEGs) (|Log2(fold change)| > 0.5, *Padj* < 0.05, merged from DEGs in comparisons of hypoxia + NTC vs normoxia + NTC, and hypoxia + HIF1αKD vs hypoxia + NTC) across all samples, representative genes labelled. (E) and (F) GSEA results of 9621 DEGs (*Padj* < 0.05) between hypoxia and normoxia NTC organoids (E), 5904 DEGs (*Padj* < 0.05) between *HIF1A-*KD and NTC hypoxic organoids (F). (G) Stabilised form of HIF1α was induced by Tet-On system under normoxia with GFP as control. (H) Immunostaining of organoids overexpressing GFP or HIF1α for 30 days under normoxia. Representative images of 2 organoid lines. (I) and (J) HIF1α overexpression under normoxia induced HIF pathway genes (I) and differentiation genes (J). The fold changes normalised to the average of GFP-overexpression organoids. Data shown as Log_2_(fold change), n = 3 biological donors. Scale bars=100 µm. For RT-qPCR, Gene expression was normalised to *ACTB*. Statistical test: two-way ANOVA with Bonferroni’s multiple comparisons test. Significance levels: *p < 0.05, **p < 0.01, ***p < 0.001, ****p < 0.0001. See also Figure S5.

To delineate the primary effects of hypoxia and HIF1α knockdown, we cultured NTC and *HIF1A*-targeting organoids under either normoxia or hypoxia for 9 days (Figure 4D). Control RT-qPCR experiments showed that even at experimental day 9, the NTC organoids upregulated airway markers and down regulated *SFTPC* under hypoxia, moreover that airway gene expression was efficiently rescued by knocking-down *HIF1A* to 5-20% of control levels (Figure S5A). We carried out bulk-RNA sequencing at day 9 for these NTC and *HIF1A*-targeting organoids using two different NTC/*HIF1A* gRNAs and three biological donors for each condition (Figure 4D). We compared NTC organoids between hypoxia and normoxia to analyse hypoxia responsive genes, and compared NTC and *HIF1A*-targeting organoids under hypoxia to identify HIF1α regulated genes (Figure 4D). Comparing NTC organoids between hypoxia and normoxia resulted in 9621 differentially expressed genes (DEGs) (*Padj* < 0.05). Gene Set Enrichment Analysis (GSEA) revealed that the DEGs induced by hypoxia were associated with processes including hypoxia responses, glycolysis, and cell-cell adhesion (Figures 4E and S5B). These included genes involved in FGF, WNT, EGF and VEGF signalling and transcription factors associated with tube morphogenesis/developmental processes, such as *KLF4*, *KLF5*, *HOPX*, *ASCL1*, and *HES1* (Figures 4D, 4E and S5B). Additionally, the NF-κB and inflammatory response pathways were activated, consistent with the elevated regulon activities in the secretory-like cell population (Figure S3A). Conversely, genes related to the cell cycle, oxidative phosphorylation, and fat metabolism were downregulated under hypoxia (Figures 4E and S5B).

Knocking down *HIF1A* under hypoxia resulted in 5904 DEGs (*Padj* < 0.05) compared with NTC organoids under hypoxia. The downregulated genes were involved in hypoxia responses, glycolysis, cell cycle, TP53 targets and the EGFR signalling pathway (Figures 4F and S5C). Suppressing *HIF1A* in hypoxia for 9 days decreased primed progenitor markers and regulons identified in the scRNA-seq dataset (*GPC3*, *WDR91*, *CPM*, *HOPX*, *FOXA1*) but not tip progenitors (*TESC*, *SOX9*) (Figure 4D). Interestingly, surfactant protein genes (*SFTPA1/2*, *SFTPC*, *SFTPD*, *SFTA3*), other AT2 markers (*LYZ*, *NAPSA*) and AT1 markers (*AQP5*, *AGER*) also decreased after *HIF1A* depletion under hypoxia (Figures 4D, 4F and S4C), suggesting HIF1α may function to maintain alveolar gene expression in hypoxia. In addition, HIF1α inhibition upregulated genes involved in oxidative phosphorylation, lipid and cholesterol metabolism (Figures 4F and S5C). Together these data show that in addition to changing cell proliferation and metabolism, hypoxia also activates lung developmental programmes, partially via HIF1α.

To determine whether inducing HIF1α alone is sufficient to drive progenitor differentiation, we used the Dox-inducible TetON system to overexpress stabilised HIF1α (Figure 4G). The stabilised HIF1α carries three point mutations on hydroxylation sites (P402A, P564A, N803A) to prevent its degradation under normoxia or suppression of transactivation functions.^60^ Stabilised HIF1α was detected in nuclei of normoxic organoids, along with widespread KRT5^+^ cells (Figure 4H). Overexpressing stabilised HIF1α under normoxia for 30 days increased *VEGF* and *GLUT1* expression compared to control GFP overexpression, though *HIF2A* also increased by 2-fold (Figure 4I). Consistently, basal cell (*KRT5*), secretory cell (*SCGB1A1*) and airway progenitor (*MUC4*) markers were activated by HIF1α overexpression, while neuroendocrine cell (*ASCL1*) and AT2 cell (*SFTPC*) markers remained unaffected (Figure 4J). We previously reported an airway differentiation medium (AWDM) to derive basal and secretory cells from lung progenitors under normoxia.^6^ Knocking down HIF1α did not affect airway differentiation under normoxia (Figure S5D), suggesting that hypoxic and chemical signalling act independently to promote airway differentiation.

### HIF2α promotes basal, but inhibits secretory, neuroendocrine, and alveolar cell fates

To determine the role of HIF2α in regulating progenitor differentiation under hypoxia, we used the CRISPRi system to knock down *HIF2A*. Inhibiting *HIF2A* for 15 days under hypoxia promoted neuroendocrine cell differentiation (*ASCL1*, *GRP*) (Figure 5A). Extended *HIF2A* suppression for 30 days limited basal cell differentiation (KRT5), but promoted expression of secretory (*SCGB3A2*) and AT2 (*SFTPC*) markers (Figure 5A). At the protein level, more SCGB3A2^+^ and ASCL1^+^ cells but fewer KRT5^+^ cells appeared in *HIF2A*-targeting organoids than NTC organoids under hypoxia (Figure 5B).

**Figure 5.**
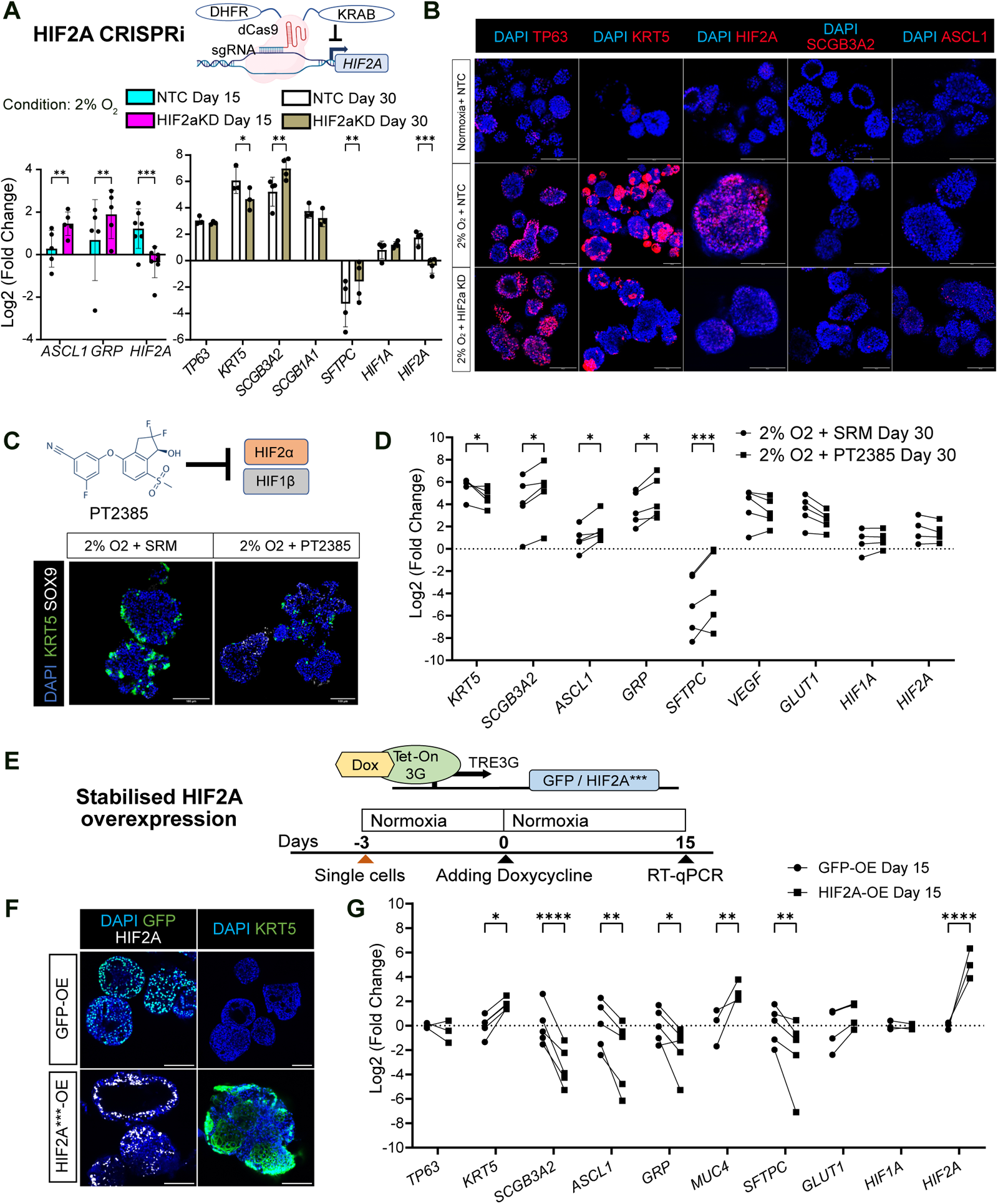
HIF2α promotes basal, but inhibits secretory, neuroendocrine, and alveolar cell fates. (A) RT-qPCR results of NTC and *HIF2A*-knock down (*HIF2A-*KD) organoids cultured under hypoxia for 15 and 30 days. The fold changes were normalised to the mean NTC + normoxia condition (not shown). Bars represent mean Log_2_(fold change) ± SD, n = 5 (day15) and 4 (day30) experimental replicates from 3 biological donors. 2 gRNAs used. (B) Immunostaining of organoids with NTC or *HIF2A-*KD induction for 30 days. Representative images of 2 organoid lines. (C) PT2385 inhibits heterodimerization between HIF2α and HIF1β. Treatment with PT2385 under hypoxia for 30 days decreased KRT5^+^ cells while increasing SOX9^+^ cells. Representative images of 2 organoid lines. (D) RT-qPCR from organoids treated with SRM only (with DMSO) or SRM + PT2385 under hypoxia for 30 days. Fold changes were normalised to the mean of normoxia + SRM condition (not shown). Data shown as Log_2_(fold change), n = 5 biological donors. (E) Stabilised form of HIF2α was induced by Tet-On system under normoxia, with GFP-overexpression as control. (F) Immunostaining of organoids overexpressing GFP or HIF2α for 15 days under normoxia. Representative images of 2 organoid lines. (G) RT-qPCR from HIF2α overexpression under normoxia for 15 days. Fold changes were normalised to the average of GFP-overexpression organoids. Data shown as Log_2_(fold change), n = 5 experimental replicates from 4 biological donors. Scale bars = 100 µm. For RT-qPCR, gene expression was normalised to *ACTB*. Statistical test: two-way ANOVA with Bonferroni’s multiple comparisons test. Significance levels: *p < 0.05, **p < 0.01, ***p < 0.001, ****p < 0.0001.

We also applied a selective HIF-2 antagonist PT2385, which inhibits HIF2α-HIF1β heterodimer formation,^61,62^ to validate the *HIF2A*-CRISPRi effects (Figure 5C). HIF2α chemical inhibition for 30 days under hypoxia reduced KRT5^+^ cells (Figure 5C), and decreased *KRT5*, but promoted secretory (*SCGB3A2*), neuroendocrine (*ASCL1*, *GRP*) and AT2 (*SFTPC*) gene expression (Figure 5D), consistent with the genetic knock-down results.

Overexpressing a stabilised form of HIF2α which carries three point mutations P405A, P531A, N847A^60^ under normoxia promoted basal cell differentiation, but inhibited secretory, neuroendocrine and AT2 marker expression (Figures 5E-G). In contrast to HIF1α promoting airway differentiation, these results consistently showed HIF2α promoted lung progenitor differentiation towards basal cells, but inhibiting AT2 and other airway cell fates.

### KLF4 and KLF5 promote basal and secretory cell fates downstream of the HIF pathway

Combining the list of HIFα-binding genes from targeted DamID-seq and hypoxia-activated genes from bulk RNA-seq, we predicted primary target genes of HIF1α and HIF2α (Figure 6A). Many development-related transcription factors were identified, such as *KLF5* in HIF1α and HIF2α common target genes (904 genes), *ASCL2* in HIF1α target genes (189 genes), and *KLF4* in HIF2α target genes (397 genes) (Figure 6A). Among them, KLF4 is known to regulate oesophageal basal cell differentiation and is directly activated by HIF1α in vascular smooth muscle cells.^63,64^ KLF5 regulates proliferation and differentiation of prostate basal cells,^65^ and is enriched in lung basal cells in a fetal lung epithelial cell dataset.^48^ We found that HIF1α and HIF2α consistently showed enriched DamID signals surrounding *KLF4* or *KLF5* (Figure 6B). As hypoxia promotes the differentiation of lung progenitors to airway cells, including basal cells, we decided to test functions of KLF4 and KLF5 during this process.

**Figure 6.**
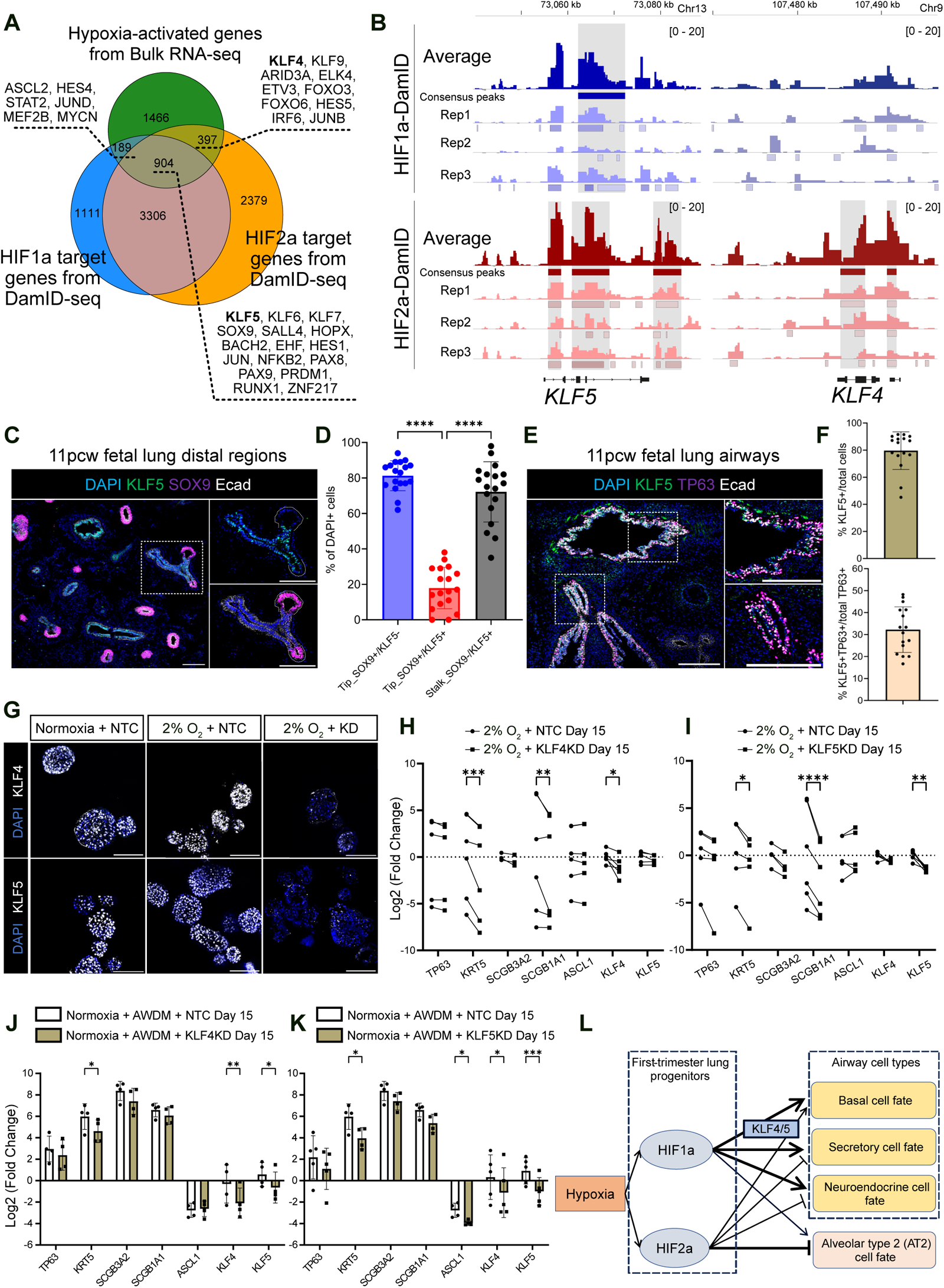
KLF4 and KLF5 promote basal and secretory cell fates downstream of the HIF pathway. (A) Venn diagram of HIF1α and HIF2α binding genes from targeted DamID-seq (FDR < 0.01) and hypoxia-activated genes from bulk RNA-seq (Log2(fold change) > 0.5, *Padj* < 0.05, DEGs of hypoxia + NTC vs normoxia + NTC). (B) Gene track views showing respective and averaged DamID signals from three biological replicates over *KLF4* and *KLF5* with consensus peaks labelled. (C-F) Immunostaining and quantification of 11 pcw human fetal lung sections showing KLF5, epithelium (Ecad), tip cells (SOX9) and differentiating basal cells (TP63) in distal regions (C, D) and airways (E, F). Quantification with lung sections from 3 donors. Statistical test: one-way ANOVA with Bonferroni’s multiple comparisons test. ****p < 0.0001. (G) Immunostaining of organoids with NTC, *KLF4*-knock down (KD) (upper panel) or *KLF5*-KD (lower panel) by CRISPRi induction for 15 days. Representative images of 2 organoid lines for each gene. (H) and (I) RT-qPCR results of *KLF4*-KD (H) and *KLF5*-KD (I) compared with NTC in SRM under hypoxia for 15 days. Data shown as Log_2_(fold change), n = 6 experimental replicates from 4 (*KLF4*), 5 (*KLF5*) biological donors. 2 gRNAs used for each gene. (J) and (K) RT-qPCR results of *KLF4*-KD (J) and *KLF5*-KD (K) compared with NTC in airway differentiation medium (AWDM) under normoxia for 15 days. Data shown as Log_2_(fold change), n = 4 (*KLF4*), 5 (*KLF5*) experimental replicates from 2 (*KLF4*), 3 (*KLF5*) biological donors. 2 gRNAs used for each gene. (L) Proposed mechanisms underlying hypoxia-induced progenitor fate decisions. Scale bars = 100 µm. RT-qPCR gene expression was normalised to *ACTB*. Statistical test: two-way ANOVA with Bonferroni’s multiple comparisons test. Significance levels: *p < 0.05, **p < 0.01, ***p < 0.001, ****p < 0.0001. See also Figure S6.

From the bulk RNA-seq data, hypoxia activated the transcription of both *KLF4* and *KLF5* (Figure S6A), while knocking down *HIF1A* specifically decreased *KLF5* expression (Figure S6B). In contrast, knocking down *HIF2A* decreased *KLF4* expression (Figure S6C). Overexpression of HIF1α or HIF2α individually under normoxia was sufficient to upregulate *KLF4*, but not *KLF5* (Figures S6D and S6E). Therefore, HIF1α and HIF2α both directly bind *KLF4* and *KLF5* but differentially regulate their expression.

In the organoid scRNA-seq dataset, *KLF4* increased expression in hypoxia-induced cell types while *KLF5* was expressed more ubiquitously (Figure S6F). In a human fetal lung atlas,^6^ however, both *KLF4* and *KLF5* have higher expression levels in airway progenitors, differentiated airway cells, late-stage tip and alveolar cells than early/mid tip cells (Figure S6G). In first-trimester human fetal lungs, the tip region, marked by SOX9, was mostly devoid of KLF5 expression (< 20% of tip cells scored as SOX9^+^KLF5^low^ cells). The level of KLF5 increased across the tip-stalk boundary and on average 70% stalk cells were KLF5^+^, coinciding with the tip-stalk progenitor state transition (Figures 6C and 6D). In differentiated airway epithelium located more proximally within the tissue, approximately 80% epithelial cells were KLF5^+^, and on average 30% TP63^+^ cells co-expressed KLF5 (Figures 6E and 6F). In contrast, KLF4 was expressed more broadly throughout the fetal lung including in mesenchymal-derived cells (Figure S6H), consistent with transcriptomic data from a fetal lung cell atlas.^6^ These data collectively suggest that KLF4 and KLF5 associate with lung epithelial differentiation, particularly of basal and secretory cells during airway differentiation.

We tested the hypothesis that KLF4 and KLF5 regulate airway differentiation downstream of HIF1α and HIF2α by using CRISPRi to inhibit *KLF4* or *KLF5* transcription separately. KLF4 and KLF5 were expressed in the NTC organoids under normoxia and hypoxia, but depleted in the CRISPRi organoids (Figure 6G). *KLF4*-CRISPRi and *KLF5*-CRISPRi both limited basal (*KRT5*) and secretory (*SCGB1A1*) cell markers but not a neuroendocrine cell marker (*ASCL1*) under hypoxia, suggesting that KLF4 and KLF5 indeed function downstream of HIFα in promoting basal and secretory cell fates (Figures 6H and 6I).

To determine whether KLF4 and KLF5 are also required for biochemical-induced airway differentiation, we cultured the *KLF4* and *KLF5*-targeting organoids in AWDM under normoxia. Consistent with hypoxia-induced airway differentiation, *KLF4* and *KLF5* inhibition resulted in lower levels of basal (*TP63*, *KRT5*) and secretory (*SCGB3A2*, *SCGB1A1*) cell differentiation during biochemical-mediated differentiation (Figures 6J and 6K). *KLF5* inhibition also decreased *ASCL1* expression in this condition. Interestingly, inhibiting either *KLF4* or *KLF5* decreased the other’s expression, suggesting a mutual regulation between these two transcription factors. These data suggest that hypoxia-induced and chemical-induced airway differentiation may converge through KLF4/KLF5 to promote basal and secretory cell fates.

Taken together, we propose a working model to explain the roles of HIF1α and HIF2α in regulating first-trimester progenitor fate decisions under hypoxia (Figure 6L). HIF1α is required for both airway differentiation and maintaining alveolar programmes, while HIF2α promotes basal cell but inhibits other airway and especially alveolar cell fates. KLF4 and KLF5 are direct HIF target genes mediating basal and secretory cell differentiation.

### Chronic hypoxia reprogrammes human alveolar type 2 cells to airway cells

We have demonstrated that chronic hypoxia promotes airway differentiation of fetal lung progenitors (Figure 1D), but prohibits their alveolar differentiation (Figure 1G). However, it remains unclear whether differentiated human AT2 cells can transdifferentiate to airway cells in response to hypoxia. We therefore established human AT2 organoids to test this hypothesis. We isolated and transduced alveolar-fated epithelial cells from human second-trimester (17-21 pcw) lung distal regions with an *SFTPC-GFP* reporter construct, sorted GFP^+^ cells, and then differentiated them in Alveolar Type 2 Medium (AT2M) to obtain fetal-derived AT2 (fdAT2) cells (Figure 7A).^66^ The fdAT2 cells recapitulate mature human AT2 cell features regarding surfactant protein production, lamellar body formation and transcriptomic profiles.^66^ We then cultured fdAT2 cells under either normoxia or hypoxia. The fdAT2 cells actively proliferated and maintained *SFTPC*-GFP expression over passages under normoxia (Figure 7B). In contrast, within 1 week of hypoxia treatment the fdAT2 cells decreased *SFTPC*-GFP levels and all lost *SFTPC*-GFP signals by 30 days (Figure 7B). In line with the decrease of the *SFTPC*-GFP signal, the fdAT2 cells gradually lost canonical AT2 markers (pro/mature-SFTPB, mature-SFTPC, ABCA3, LAMP3), reduced proliferation (Ki67) and acquired basal cell markers (TP63, KRT5) as confirmed by immunostaining (Figure 7C).

**Figure 7.**
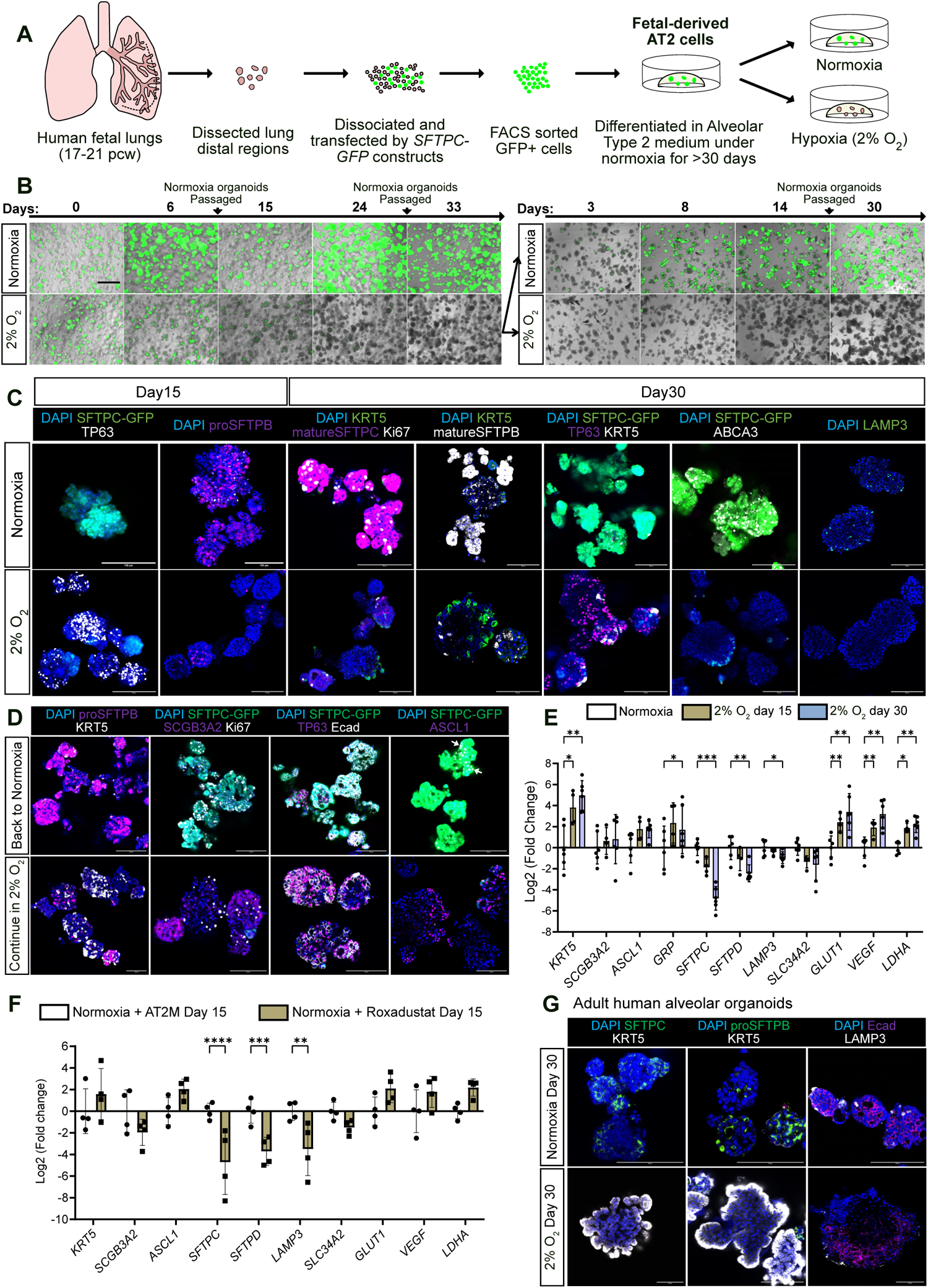
Chronic hypoxia reprogrammes human alveolar type 2 cells to airway cells. (A) Experiment design. The distal epithelial cells were isolated from second-trimester human fetal (17-21 pcw) lungs, transduced by *SFTPC-GFP* reporter, FACS sorted, and cultured in Alveolar Type 2 Medium as fdAT2 organoids under normoxia or hypoxia. (B) Microscope images of merged brightfield and GFP channels. The fdAT2 organoids with *SFTPC-GFP* reporter were cultured under normoxia (with passaging) or hypoxia (without passaging) for 33 days. The hypoxia-treated organoids were split and cultured under normoxia or hypoxia for another 30 days. Scale bars = 600 µm. Representative images of 2 organoid lines. (C) Immunostaining of AT2 markers (SFTPC, SFTPB, ABCA3, LAMP3), basal cell markers (TP63, KRT5), and proliferation marker (Ki67) of fdAT2 organoids cultured under normoxia or hypoxia for 15 and 30 days. Representative images of 3 organoid lines. Scale bars = 100 µm. (D) Immunostaining of hypoxia pre-treated fdAT2 organoids cultured under normoxia or hypoxia for 30 days. Representative images of 2 organoid lines. Arrows indicate *SFTPC-*GFP^+^ASCL1^+^ cells. Scale bars = 100 µm. (E) RT-qPCR of fdAT2 organoids cultured under normoxia or hypoxia for 15 and 30 days. Fold changes were normalised to the mean of the normoxia condition. Data shown as mean Log_2_(fold change) ± SD, n = 6 experimental replicates from 4 biological donors. Statistical test: two-way ANOVA with Dunnett’s multiple comparisons test. (F) RT-qPCR of fdAT2 organoids treated with Roxadustat under normoxia for 15 days. Fold changes were normalised to the mean of fdAT2 without Roxadustat treatment (with DMSO). Data shown as mean Log_2_(fold change) ± SD, n = 4 experimental replicates from 3 biological donors. (G) Immunostaining of adult AT2 organoids cultured under normoxia or hypoxia for 30 days. Representative images of 2 organoid lines. Scale bars = 100 µm. Gene expression was normalised to *ACTB* in RT-qPCR. Significance levels: *p < 0.05, **p < 0.01, ***p < 0.001.

To determine whether this hypoxia-induced alveolar cell reprogramming is reversible, we split the day 33 hypoxia-treated fdAT2 organoids into either normoxia or hypoxia conditions and cultured for a further 30 days. Many organoids switched on *SFTPC*-GFP within 3 days after returning to normoxia, and the *SFTPC*-GFP^+^ cell proportion increased over 1-month of culture (Figure 7B). Moreover, upon returning to normoxia the organoids regained pro-SFTPB, recovered proliferation (Ki67), and mostly lost basal cell markers (TP63, KRT5) though some cells retained the neuroendocrine marker (ASCL1) (Figure 7D). In comparison, the organoids cultured under hypoxia for 2 months remained negative for AT2 markers (pro-SFTPB) and positive for airway markers (TP63, KRT5, SCGB3A2, ASCL1) (Figure 7D).

To monitor gene expression changes in the fdAT2 organoids under hypoxia, we performed RT-qPCR. The fdAT2 organoids downregulated alveolar markers (*SFTPC*, *SFTPB*, *LAMP3*, *SLC34A2*) but upregulated basal cell marker (*KRT5*) and HIF target genes (*GLUT1*, *VEGF*, *LDHA*) in a time-dependent way (Figure 7E). The neuroendocrine markers (*ASCL1*, *GRP*) were also moderately activated. Furthermore, stabilising the HIF pathway in normoxia by adding Roxadustat could recapitulate the hypoxia effects in the fdAT2 organoids (Figure 7F). These findings suggest that chronic hypoxia directly reprogrammed human AT2 cells into airway cells through the HIF pathway.

The fdAT2 organoids have been shown to closely resemble adult AT2 cells in many aspects.^66^ However, to confirm that the hypoxia response is maintained in adult AT2 cells, we also isolated distal epithelial cells from adult human lung parenchyma and cultured them in the serum-free feeder-free (SFFF) medium as adult-derived AT2 (adAT2) organoids as previously reported.^67^ The adAT2 organoids expressed SFTPC, SFTPB and LAMP3 under normoxia, but lost alveolar markers and strongly expressed KRT5 after 30-day hypoxia culture, consistent with the fdAT2 organoid response (Figure 7G).

## DISCUSSION

Hypoxia prevails during early human development and occurs in adult pathological states. Although lungs are the adult site of gas exchange, the embryonic and early pseudoglandular (airway formation) stages of human lung development happen in a hypoxic environment. Injuries and chronic diseases, such as influenza infection and fibrosis, can also lead to local hypoxia in adult lungs. Here we show that the first-trimester lung progenitors directly respond to hypoxia and autonomously differentiate to airway cells in the self-renewing culture condition at the expense of alveolar gene expression. Activation of the HIF pathway under normoxia phenocopied hypoxia effects. HIF1α and HIF2α have largely overlapping target genes, but distinctly control this differentiation process. Furthermore, hypoxia exposure also efficiently converts differentiated AT2 cells to airway cells.

Through single cell transcriptomics, we found that tip and primed progenitors co-exist in the self-renewing organoids. The tip progenitors slowed proliferation, but didn’t actively differentiate under hypoxia. In contrast, the primed progenitors pre-activated developmental genes and readily gave rise to multiple types of airway cells (basal, neuroendocrine, secretory-like, and hillock-like cells) under hypoxia. We also noticed a small fraction of differentiating cells and HRE-ODD-GFP activation in organoids cultured under normoxia, which may result from local hypoxia as organoid size and density increase. Frequent passaging prevented the accumulation of hypoxia effects during routine organoid maintenance.

Here we found that HIF1α and HIF2α both promoted basal cell fate, but had opposing effects on secretory, neuroendocrine, and alveolar fates. Specifically, HIF1α was required for neuroendocrine cell differentiation and maintaining expression of surfactant protein genes under hypoxia, while HIF2α inhibited both lineages. Consistent with this observation, during mouse tracheal repair HIF1α was observed to promote, while HIF2α inhibited, the differentiation of basal cells to neuroendocrine cells.^24^ In newborn rodents, HIF1α is required for alveolar development by supporting normal vascularization and surfactant production.^68–70^ In contrast, inhibition of HIF2α facilitates alveolar regeneration.^32^ Corresponding to their functional differences, HIF1α and HIF2α have distinct expression patterns in human fetal lungs,^71^ and regulate partially overlapping target genes in cell lines.^72,73^ Through targeted DamID-seq, we found that HIF1α and HIF2α shared the majority of but not identical target genes in the lung epithelial progenitors. HIF1α and HIF2α bound at transcription factors governing differentiation towards basal and secretory cells (KLF4, KLF5), suggesting that HIF1α and HIF2α regulate airway fate decisions by promoting the expression of core airway differentiation transcription factors.

Hypoxia is prevalent in chronic lung diseases, such as lung fibrosis. Fibrotic lungs frequently show bronchiolization of the alveolar regions. Aberrant basal cell emergence can be a HIF-dependent process. For example, after severe influenza damage HIF1α drives basal-like cell expansion from lineage-negative progenitors in mouse distal lungs.^29–31^ HIF1α also regulates the conversion of mouse AT2 cells to transient intermediate cells (known as DAPT - damage associated transient progenitors, or PATS - pre-alveolar type-1 transitional cell state) which can aberrantly accumulate in fibrotic lungs.^74–76^ Moreover, HIF2α inhibition attenuates mouse lung fibrosis after repetitive bleomycin injuries, promoting alveolar regeneration.^32^ Pathogenic mesenchyme has also been shown to promote transdifferentiation of human AT2 cells into KRT5^+^ basal cells in IPF (Idiopathic Pulmonary Fibrosis) lungs.^35^ Our observations indicate that human AT2 cells can directly sense hypoxia and autonomously differentiate to airway cells. The pathogenic niche and hypoxic environment may thus together promote disease progression.

## ACKNOWLEDGEMENTS

We acknowledge the Gurdon Institute Imaging Facility, Bioinformatics group, and Animal Facility. We thank Dr. Kyungtae Lim for sharing fetal-derived AT2 cells. Z.D. is supported by a Wellcome Trust PhD studentship (222275/Z/20/Z). E.L.R. is supported by the Medical Research Council (MRC) (MR/P009581/1; MR/S035907/1). J.v.d.A. is supported by a Wellcome Clinical Research Career Development Fellowship (219615/Z/19/Z), a Wellcome Discovery Award (226653/Z/22/Z), a UKRI BBSRC Responsive Mode Research Grant (BB/X00256X/1) and acknowledges core funding from the MRC to the MRC Mitochondrial Biology Unit (MC_UU_00028/8). J.A.N. is supported by a Wellcome Senior Clinical Research Fellowship (215477/Z/19/Z) and a Lister Institute Research Fellowship. Core funding to the Gurdon Institute from the Wellcome Trust (203144/Z/16/Z) and CRUK (C6946/A24843).

## AUTHOR CONTRIBUTIONS

Z.D., J.A.N. and E.L.R. conceptualised the project. Z.D. designed and performed most experiments and analyses. N.W. analysed DamID-seq data. A.A. assisted RT-qPCR and immunohistochemistry experiments of KLF5. D.D. and Z.D. prepared sequencing libraries for DamID-seq. A.J.R. supervised scRNA-seq and bulk RNA-seq analyses. J.v.d.A. and J.A.N. provided resources. J.v.d.A., J.A.N. and E.L.R. acquired funding and supervised the project. Z.D. wrote the original manuscript. Z.D. and E.L.R. edited the manuscript with input from all authors.

## MATERIALS AND METHODS

### Human fetal and adult lung tissue

Human embryonic and fetal lung tissues were provided from Cambridge University Hospitals NHS Foundation Trust under NHS Research Ethical Committee (96/085) and the MRC/Wellcome Trust Human Developmental Biology Resource (London and Newcastle, University College London (UCL) site REC reference: 18/LO/0822; Newcastle site REC reference: 18/NE/0290; Project 200454; www.hdbr.org). Stages of the samples were evaluated by external appearance and measurements to determine their age in post-conception weeks (pcw). Human adult lung tissues were provided from Cambridge Biorepository for Translational Medicine (CBTM) (reference: 15/EE/0152). None of the samples used for this study had known genetic abnormalities.

### Mouse breeding

Mice were bred and maintained under specific-pathogen-free conditions at the Gurdon Institute of the University of Cambridge. All mouse procedures were approved by the University of Cambridge Animal Welfare and Ethical Review Body and carried out under a UK Home Office License (PPL: PEEE9B8E4) in accordance with the Animals (Scientific Procedures) Act 1986.

### Derivation and maintenance of human fetal lung epithelial progenitor organoids

Human fetal lung epithelial progenitor organoids were derived as previously reported.^4^ Briefly, human fetal lung tissues (7-9 pcw) were dissociated with Dispase (8 U/mL Thermo Fisher Scientific, 17105041) at room temperature for 2 min. Mesenchyme was removed by forceps. Branching epithelial tips were micro-dissected, transferred into basement membrane extract (BME, Bio-Techne, 3533-010-02) on 24-well suspension culture plates (M9312-100EA, Greiner). The organoids were expanded in Self-Renewal Media (SRM) consisting of AdvDMEM+++ medium [Advanced DMEM/F12 (ThermoFisher Scientific, 12634010) with 1x GlutaMax (ThermoFisher Scientific, 35050061), 10 mM HEPES (ThermoFisher Scientific, 15630056) and 100 U/mL Penicillin/Streptomycin (ThermoFisher Scientific, 15140122)] and supplements [N2 (1:100, ThermoFisher Scientific, 17502–048), B27 (1:50, ThermoFisher Scientific, 12587–010), 1.25 mM N-acetylcysteine (Merck, A9165), 5% v/v R-spondin condition medium (Stem Cell Institute Tissue Culture, University of Cambridge), 50 ng/mL recombinant human EGF (PeproTech, AF-100-15), 100 ng/mL recombinant human Noggin (PeproTech, 120-10C), 100 ng/mL recombinant human FGF10 (PeproTech, 100-26), 100 ng/mL recombinant human FGF7 (PeproTech, 100-19), 3 μM CHIR99021 (Stem Cell Institute Tissue Culture, University of Cambridge) and 10 μM SB431542 (Bio-Techne, 1614)] in a CO_2_ incubator (20-21% O_2_, 5% CO_2_) or hypoxia incubator (2-5% O_2_, 5% CO_2_). Organoids cultured under normoxia were passaged every 5-7 days depending on the confluence. For passaging organoids, fresh cold (4℃) AdvDMEM+++ was used to disrupt the BME mechanically and harvest organoids. The organoids were pelleted by centrifugation and dissociated using TrypLE (Thermo Fisher Scientific, 12605010) at 37℃ for 10 min, or sheared by pipetting. The cells or organoid pieces were washed in AdvDMEM+++ and resuspended in BME according to subculture ratios. SRM was supplemented with 10 µM Y-27632 for first 3 days. Organoids cultured under hypoxia were passaged by mechanical shearing using a 200 µL pipette tip around every 2 weeks. All fetal lung organoids were detected negative of mycoplasma.

### Molecular cloning

For mutated HIF1α and HIF2α overexpression, the *HIF1A* and *HIF2A* CDS were cloned from plasmids gifted from William Kaelin (Addgene, #87261, #25956) and inserted into Tet-ON vectors with EF1a-TagRFP-2A-tet3G.^58,60^ For CRISPRi, the gRNA sequences targeting *HIF1A*, *HIF2A*, *KLF4*, *KLF5* were selected from published dataset and inserted into U6-gRNA-EF1a-EGFP-CAAX lentiviral vectors (Addgene, #167936).^58,59^ For targeted DamID, the wild-type *HIF1A* and *HIF2A* CDS were inserted into DamID vectors with SFFV-mNeonGreen as upstream open reading frame.^43^ The tetON-KRAB-dCas9-DHFR-EF1a-TagRFP-2A-tet3G plasmid (Addgene, #167935), NTC (non-targeting control) plasmid, *SFTPC-GFP* reporter plasmid and HRE-ODD-GFP reporter plasmid were as previously described.^40,57,58^

### Lentiviral production and organoid transduction

HEK293T cells were grown in 10-cm dishes to 80% confluency before transfection with the lentiviral vector (10 μg) with packaging vectors including pMD2.G (3 μg, Addgene, # 12259), psPAX2 (6 μg, Addgene, #12260) and pAdVAntage (3 μg, Promega, E1711) using Lipofectamine 2000 Transfection Reagent (Thermo Fisher Scientific, 11668019) according to manufacturer’s protocol. After 16 hrs, medium was refreshed. Supernatant containing lentivirus was harvested at 24 hrs and 48 hrs after medium refreshing and pooled together. Supernatant was centrifuged to remove cell fragments and passed through 0.45 μm filter. Lentivirus containing > 10 kb length insert (inducible CRISPRi) was concentrated using AVANTI J-30I centrifuge (Beckman Coulter) with JS-24.38 swing rotor at 100,000g for two hours at 4°C and pellets were dissolved in 200 μL PBS. Alternatively, the lentivirus was concentrated using Lenti-X™ Concentrator (Takara, 631232) following the manufacturer’s protocol. For transduction, the organoids were dissociated by TrypLE and cultured in SRM with 10 µM Y-27632 in suspension with packaged viruses (10 μL viruses for each confluent-well organoids in 24-well plates) overnight. The cells were washed by AdvDMEM+++ and cultured in SRM with 10 µM Y-27632 for first 3 days. After 5-7 days, the organoids were dissociated for cell sorting (Sony SH800Z Cell Sorter) with wild-type cells as the negative control. For Tet-ON overexpression, 2 µg/mL Dox (Merck, D9891) was added into SRM. For CRISPRi, 2 µg/mL Dox and 10 µM TMP (Merck, 92131) were used.

### Derivation and maintenance of human fetal-derived AT2 (fdAT2) cells

The dissection and isolation of distal epithelial cells from human second-trimester (17-21 pcw) lungs were as previously described.^40,66^ Briefly, the lung distal regions were cut into small pieces and dissociated in 5 mL of enzyme mixture (0.125 mg/mL Collagenase, Merck, C9891; 1 U/mL Dispase, Thermo Fisher Scientific, 17105041; 10 U/mL DNAase, Merck, D4527) at 37°C for 1 hour with rotation. The cells were washed with AdvDMEM+++ medium and filtered through a 40 μm strainer. The supernatant was removed after centrifugation and the cell pellet was resuspended in red blood cell lysis buffer (BioLegend, 420301) for 5 min, and washed with AdvDMEM+++ medium. The dissociated cells were enriched for epithelial cells by Magnetic-activated cell sorting (MACS) (buffer: 1x PBS, 1% BSA, and 2 mM EDTA) with CD326 (EpCAM) microbeads (Miltenyi Biotec, 130-061-101) according to the manufacturer’s instructions. The enriched cells were resuspended in BME and seeded into multi-well plates for culture with the Alveolar Type 2 Medium (AT2M) [AdvDMEM+++, 1X B27 supplement (without Vitamin A), 1x N2 supplement, 1.25 mM n-Acetylcysteine, 10 mM CHIR99021, 50 μM Dexamethasone (Merck, D4902), 10 µM Y-27632, 0.1 M 8-Bromoadenosine 3’5’-cyclic monophosphate (cAMP; Merck, B5386), 0.1 M 3-Isobutyl-1-methylxanthine (IBMX; Merck, 15679), 50 mM DAPT (Merck, D5942), and 10 mM A83-01 (Tocris, 2939)]. Medium was changed every 3 days and the organoids were passaged around every 2 weeks. Alternatively, the dissociated cells were transduced by the *SFTPC-GFP* reporter lentiviral construct in suspension in AT2M overnight. The cells were collected and expanded in BME in AT2M for 5-6 days. The SFTPC-GFP+ cells were sorted (Sony SH800Z Cell Sorter) and cultured in AT2M. All fdAT2 lung organoids were detected negative of mycoplasma.

### Human adult AT2 cell isolation and culture

Human adult lung parenchyma was dissected and dissociated as previously described.^67^ Briefly, human distal lung edges were cut into small pieces, and digested with 10 mL of enzyme mixture (Collagenase: 1.68 mg/mL, Dispase: 5 U/mL, DNase: 10 U/mL) at 37°C for 2-3h with rotation and pipetting in the middle to assist digestion. The cells were washed with AdvDMEM+++ medium and filtered through a 40μm strainer. The supernatant was removed after centrifugation at 500 g for 5 min and the cell pellet was resuspended in red blood cell lysis buffer (BioLegend, 420301) for 10 min, and washed with AdvDMEM+++ medium. Total cells were centrifuged at 500 g for 5 min and the cell pellet was processed for AT2 isolation by Magnetic-activated cell sorting (MACS) with CD326 (EpCAM) microbeads (Miltenyi Biotec, 130-061-101) according to the manufacturer’s instructions. CD326 selected cells were resuspended in BME (5-10k cells per 30 uL BME drop) and seeded into multi-well plates for culture with serum-free feeder-free (SFFF) medium as previously reported.^67^ Medium was changed every 3 days and the organoids were passaged every 2-3 weeks. All adult lung organoids were detected negative of mycoplasma.

### Airway and alveolar differentiation of first-trimester lung progenitors

After growing in SRM from single cells for 3 days, the first-trimester epithelial progenitor organoids were cultured in the Airway Differentiation Medium (AdvDMEM+++, 1X B27, 1X N2, 1.25 mM N-acetylcysteine, 100 ng/mL FGF10, 100 ng/mL FGF7, 50 nM Dexamethasone, 0.1 mM cAMP, 0.1 mM IBMX, 10 µM Y-27632), or the Alveolar Differentiation Medium (AdvDMEM+++, 1X B27, 1X N2, 1.25 mM N-acetylcysteine, 10 mM CHIR99021, 50 mM DAPT, 10 μM SB431542, 50 nM Dexamethasone, 0.1 mM cAMP, 0.1 mM IBMX, 10 µM Y-27632) as previously described.^6,40^ Medium was changed every 3 days and organoids were differentiated for 9-15 days.

### Immunohistochemistry

Human embryonic and fetal lungs were fixed at 4°C overnight in 4% (w/v) paraformaldehyde in PBS. Fixed lungs were washed in 15%, 20% and 30% (w/v) sucrose in PBS at 4°C for 1 hour and incubated in 1:1 (v/v) mixture of optimal cutting temperature compound (OCT, Tissue-tek, 4583):30% sucrose (in PBS) at 4°C overnight. The lungs were finally embedded and frozen in 100% OCT and stored at −70°C before sectioning. For immunostaining, fetal lung cryosections (10 µm) were washed in PBS and incubated in PBS with 0.3% Triton X-100 (0.3% PBTX) for 10 minutes. The sections were incubated in blocking buffer (1% bovine serum albumin, 5% normal donkey serum in 0.3% PBTX) at room temperature for 1 hour and incubated with primary antibodies (KLF4, 1:500, Proteintech, 11880-1-AP; KLF5, 1:500, Proteintech, 21017-1-AP; SOX9, 1:600, Merck, AB5535; TP63, 1:600, Cell Signaling Technology, 13109; E-cadherin, 1:1000, Thermo Fisher Scientific, 13-1900) at 4℃ overnight. The sections were washed in PBS and incubated with secondary antibodies (donkey anti-rabbit 488, 1:1000, Invitrogen, A-21206; donkey anti-goat 594, 1:1000, Invitrogen, A-11058; donkey anti-rat 647, 1:1000, Jackson Immunoresearch, 712-605-153) at room temperature for 2 hours. The sections were stained with DAPI (1 µg/mL) at room temperature for 20 minutes, washed and mounted in Fluoromount (Sigma) for imaging by Leica SP8 confocal microscope and Nikon AxR confocal microscope. Images were processed using Fiji (version 2.15.1).

### Organoid whole-mount immunostaining

The organoids were released from BME by washing in cold (4℃) AdvDMEM+++ medium and fixed in 4% PFA on ice for 30min. The organoids were then washed in PBS 3 times and incubated in 0.3% PBTX for 1 hour at 4℃. The organoids were blocked at 4℃ overnight, followed by primary antibody incubation (SOX2, 1:500, Bio-techne, AF2018; SOX9, 1:500, Merck, AB5535; SOX9, 1:500, R&D Systems, AF3075; TP63, 1:400, Cell Signaling Technology, 13109; TP63, 1:400, R&D Systems, AF1916; KRT5, 1:500, BioLegend, 905901; SCGB3A2, 1:800, Abcam, ab181853; SCGB1A1, 1:800, Proteintech, 10490-1-AP; E-cadherin, 1:1000, Thermo Fisher Scientific, 13-1900; Fibronectin, R&D Systems; NKX2.1, 1:500, Abcam, ab76013; proSFTPC, 1:400, Merck, AB3786; ZO1, 1:400, Invitrogen, 40-2200; ASCL1, 1:400, Abcam, EPR19840; Ki67, 1:500, Invitrogen, 14-5698-82; Laminin, 1:500, Abcam, ab11575; KRT13, 1:500, Abcam, ab79279; HIF1α, 1:300, Novus Biologicals, NB100-134; HIF2α, 1:300, Novus Biologicals, NB100-122; KLF4, 1:400, Proteintech, 11880-1-AP; KLF5, 1:400, Proteintech, 21017-1-AP) at 4℃ overnight. The organoids were washed in PBS and incubated in secondary antibodies (donkey anti-chick Alexa 488, 1:1000, Jackson Immune, 703-545-155; donkey anti-rabbit 488, 1:1000, Invitrogen, A-21206; donkey anti-mouse 488, 1:1000, Invitrogen, A-21202; donkey anti-rat 488, 1:1000, Invitrogen, A-21208; donkey anti-mouse 594, 1:1000, Invitrogen, A-21203; donkey anti-rabbit 594, 1:1000, Invitrogen, A-21207; donkey anti-goat 594, 1:1000, Invitrogen, A-11058; donkey anti-sheep 594, 1:1000, Jackson Immunoresearch, 713-585-147; donkey anti-rat 647, 1:1000, Jackson Immunoresearch, 712-605-153; donkey anti-rabbit 647, 1:1000, Invitrogen, A-31573; donkey anti-mouse 647, 1:1000, Invitrogen, A-31571; donkey anti-goat 647, 1:1000, Invitrogen, A-21447) at 4℃ overnight. After DAPI staining (1 µg/mL) at 4℃ for 1 hour, the organoids were processed through a thiodiethanol series (25%, 50%, 75% and 97% v/v concentration in PBS) at 4℃ followed by mounting in 97% thiodiethanol and imaging on Leica SP8 or Nikon AxR confocal microscopes. Images were processed using Fiji (version 2.15.1).

### Western blot

The organoid samples were harvested, lysed with RIPA buffer (Merck, R0278) after removing BME, and then run on 12.5% SDS-PAGE gels. Proteins were transferred onto PVDF membranes with BioRad Mini Trans-Blot system (BioRad, Mini Trans-Blot® Cell). The membranes were blocked with 5% skimmed milk in 0.1% Tween-20/TBS (TBST) for 30 minutes at room temperature, and incubated at 4°C overnight with primary antibodies (HIF1α, 1:1000, Novus Biologicals, NB100-134; HIF2α, 1:1000, Cell Signaling Technology, 7096; β-Actin, 1:5000, Merck, A1978) in 0.1% skimmed milk in TBST buffer (blocking buffer). After washing with TBST, the membranes were incubated with secondary antibodies conjugated with fluorescence dyes (anti-mouse IRDye® 800CW, 1:5000, Abcam, ab216774; anti-rabbit IRDye® 680RD, 1:5000, Abcam, ab216779) at room temperature for 3 hours. The membranes were washed with TBST and developed using the Li-Cor Odyssey imaging system.

### RNA extraction, reverse transcription and RT-qPCR analysis

Organoids were harvested and the RNA was extracted using RNeasy Plus Mini Kit (Qiagen, 74134). The cDNA was synthesized using High-Capacity cDNA Reverse Transcription Kit (Thermo Fisher Scientific, 4368814). Incubation at 25 ℃ for 10 minutes, 37 ℃ for 2 hours and 85 ℃ for 5 minutes. For RT-qPCR, diluted cDNA was mixed with primers and PowerUp SYBR Green Master Mix (Thermo Fisher Scientific, A25741). Fold changes of target gene expression were determined by ΔΔCT methods with *ACTB* as reference gene. The data was analysed in GraphPad Prism 10 with one or two-way ANOVA with Tukey/Bonferroni/Dunnett multiple comparison tests or linear regression as stated in each figure. Significance levels: *p < 0.05, **p < 0.01, ***p < 0.001, ****p < 0.0001.

### Bulk RNA-sequencing and analysis

The extracted RNA quality was analysed with High Sensitivity RNA ScreenTape (Agilent, 5067-5579) on Agilent 4200 Tapestation. The mRNA-sequencing library preparation and sequencing were completed by Novogene (UK) Company Limited with NovaSeq 6000. 20-50 M PE150 reads were sequenced for each sample. The sequencing data was analysed with nf-core/rnaseq pipeline (version 3.9) with default settings.^77^ The reads were mapped to human genome GRCh38.p13 and quantified by STAR v2.7.10a and Salmon v1.5.2.^78,79^ The output gene count matrix was used for differential gene expression analysis with DESeq2.^80^ The differentially expressed genes (DEGs) were extracted by the contrast function by comparing hypoxia + NTC and normoxia + NTC, and hypoxia + HIF1α-knock down and hypoxia + NTC conditions separately. The DEGs (*Padj* < 0.05) were used for Gene Set Enrichment Analysis (GSEA) with Molecular Signatures Database (v2022.1.Hs).^81,82^ The heatmap was made with pheatmap package (Version 1.0.12).^83^

### Organoid single-cell RNA sequencing and analysis

Lung progenitor organoids from two fetal lungs (9 pcw) cultured under normoxia and 8, 16, 24, and 32 days of hypoxia were harvested in parallel at each time point. The organoids were dissociated into single cells using TrypLE, filtered through a 40 μm filter to achieve > 90% single cells and evaluated by Trypan Blue Solution (Thermo Fisher Scientific, 15250061) to confirm > 90% cell viability. The cells were fixed and frozen using Evercode Cell Fixation kit (Parse Biosciences). All the samples were processed together with Evercode Whole Transcriptome v2 kit (Parse Biosciences) to generate sequencing libraries. The libraries were multiplexed and sequenced by BGI Group in one T7 lane to achieve an average of 63,000 raw PE reads per cell of estimated 82,391 total cells. The sequencing data was initially processed with split-pipe (Version 1.1.1, Parse Biosciences) to combine sublibraries. The reads were mapped to human genome GRCh38.p14. The gene count matrix was used for downstream analysis in Seurat (Version 5).^42^ The cells were filtered based on gene counts 2000-7000, transcript counts > 4000, mitochondrial gene percentages < 5% and genes detected in > 100 cells to yield total 65,475 cells. Data was normalised (normalization.method = “LogNormalize”, scale.factor = 10000) and scaled with default settings. The linear dimensional reduction was based on top 2000 highly variable features. The cell clusters (using top 20 PCs, 13 neighbours, and resolution at 0.5) were curated and annotated based on canonical *in vivo* cell type markers as previously described to generate 11 cell types.^6,47^ The differentially expressed genes for each cell type were found using FindMarkers function using default parameters.. The trajectory analysis was complemented with Monocle 3 (for both partitions containing tip and primed progenitors) and Slingshot (only the major partition containing primed progenitors) by setting the root at cycling cells.^55,56^ The cell cycle scoring was calculated with CellCycleScoring function in Seurat. For mapping the organoid data to a fetal lung epithelial cell atlas,^6^ the fetal lung epithelial cells were re-clustered with Seurat default settings (except dims = 1:50 in FindNeighbors) as the reference, and the organoid data projected onto the reference UMAP structure with FindTransferAnchors, TransferData, AddMetaData and MapQuery functions. The plots were created with DimPlot, FeaturePlot, DotPlot, and RidgePlot functions. The regulons were analysed with SCENIC on downsampled organoid data (500 cells for each of the 13 original clusters) in R.^84^ The DEGs between primed progenitors and normoxic tip progenitors, and between hypoxic tip progenitors and normoxic tip progenitors, were analysed with AggregateExpression pseudobulk function and FindMarkers function. The volcano plots were generated with EnhancedVolcano package.^85^ The connect plots were based on GSEA results for the DEGs.

### Mouse embryonic lung dissection and tip progenitor organoid culture

The first day a vaginal plug was detected was designated as embryonic (E) day 0.5. The lungs of E11.5-E14.5 C57BL/6J mouse embryos were dissected. The lung buds were cut and briefly treated with Dispase I (8 U/mL) to separate the mesenchyme. The epithelial tips were seeded in BME and cultured in the mouse lung tip progenitor medium.^41^ The organoids were passaged every 5-7 days using the same approaches as for human lung organoids.

### Targeted DamID-sequencing sample preparation

The HIF1α, HIF2α and empty DamID-only lentiviral vectors were transduced to dissociated lung progenitor organoids from 3 donors as described above. 20-40% cells were transduced as checked by the mNeonGreen signals. The cells were cultured in SRM with 10 µM Y-27632 under normoxia for 3 days, and then treated with 2% O_2_ in SRM for 6 days. Medium was changed every 3 days. Then the organoids were harvested and processed for Illumina sequencing with an adapted TruSeq protocol as previously described.^86^ All samples were multiplexed and the sequencing was performed by the Cancer Research UK Cambridge Institute genomics facility using 1 lane of Illumina NovaSeq X as PE50 reads.

### Targeted DamID-sequencing data analysis

Targeted DamID-sequencing data was processed using a publicly available Snakemake workflow.^87^ Briefly, reads were first quality trimmed using TrimGalore v0.6.10 to remove adapter sequences, after which the quality was investigated using FastQC v0.12.1 and MultiQC v1.19.^88^ These trimmed reads were then aligned to the human genome (Ensembl GRCh38.110) using bowtie2 v2.5.3.^89^ To prevent signals originating from the expression vectors of *Dam-HIF1A/HIF2A* fusion genes from obscuring the analysis, the bowtie2 index was built with a FASTA file where the genome sequences of *HIF1A* and *HIF2A* were masked using a custom Python script. Subsequently, bedGraph files were generated with reads binned into fragments based on 5’-GATC-3’ sites and normalised to a separate Dam-only control sample of the same organoid line. The alignment and bedGraph generation steps were performed using damidseq_pipeline v1.5.3.^90^ The width of bins to use for mapping reads was set at 300. HIF1α and HIF2α bedGraph files from one organoid cell line (i.e. biological replicate) were quantile normalised against all the other organoids other using a custom Python script. For the purpose of visualisation of individual loci only, the logarithmic values in the bedGraph files were back-transformed. Average signal at individual loci was plotted with pyGenomeTracks v3.8.^91^ To generate bigWig files, bedGraph files were first converted to bigWig files with UCSC bedGraphToBigWig v445, then average Wig files were obtained with wiggletools v1.2.11 that were converted back to bigWig with UCSC wigToBigWig v447.^92,93^ Broad peak calling was performed with the MACS2 v2.2.9.1 subcommand callpeak (broad-cutoff = 0.1 and q = 0.05) using bam files generated by damidseq_pipeline.^94^ Dam-HIF1α/ HIF2α served as treatment samples and the Dam-only as control sample. Consensus peaks were identified only if peaks occurred in all three biological replicates with at least 1 bp overlap. These consensus peaks were obtained with bedtools multiinter v2.31.1.^95^ Consensus peaks smaller than 100 bp were extended by 100 bp on both the 5’ and 3’ end. Consensus peaks were annotated to the nearest transcription start site with the ChIPseeker v1.38.0 R package to find HIF1α and HIF2α target genes.^96^ Venn diagrams were made using the Eulerr R package using the annotated gene names corresponding to HIF1α and HIF2α consensus peaks.^97^ Profile plots for gene groups in the Venn diagrams were generated with deepTools v3.5.4.^98^ The gene ontology analysis for HIF1α and HIF2α common target genes was performed with Enrichr.^99^

---

**Figure S1.**
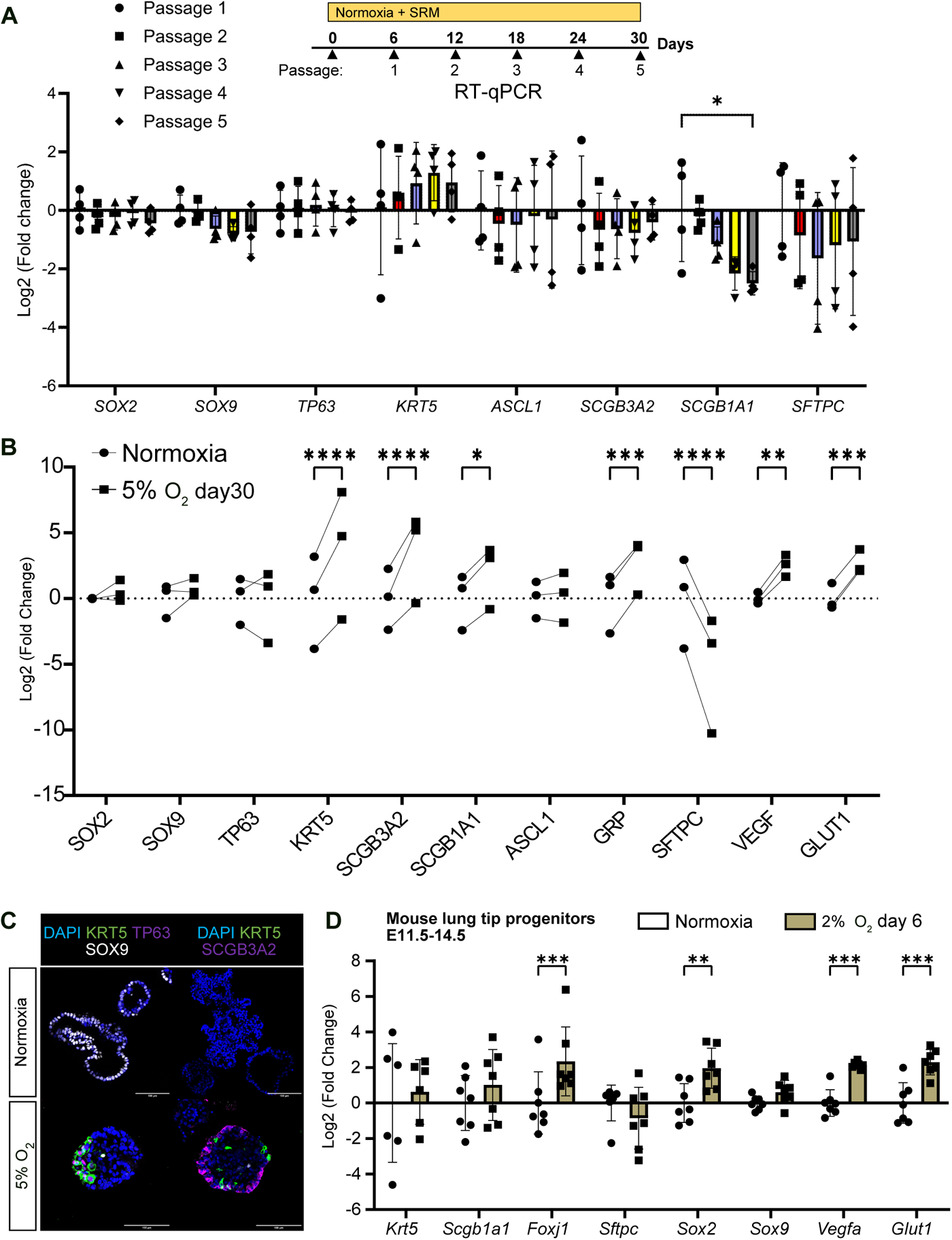
Hypoxia culture of human and mouse lung epithelial progenitors. Related to Figure 1. (A) RT-qPCR of human lung progenitor organoids cultured under normoxia with routine passaging. Fold changes were normalised to the mean of Passage 1 organoids. Bar represents mean Log_2_(fold change) ± SD, n = 4 biological donors. Statistical test: two-way ANOVA with Tukey’s multiple comparisons test. (B) RT-qPCR of human lung progenitor organoids cultured under normoxia or 5% O_2_ for 30 days. Fold changes were normalised to the mean of the normoxia condition. Data shown as Log_2_(fold change), n = 3 biological donors. Statistical test: two-way ANOVA with Bonferroni’s multiple comparisons test. (C) Immunostaining of human lung progenitor organoids cultured under normoxia or 5% O_2_ for 30 days. Representative images of 2 organoid lines. Scale bars = 100 µm. (D) RT-qPCR of mouse lung progenitor organoids derived from E11.5-14.5 embryos cultured under normoxia or 2% O_2_ for 6 days. Fold changes were normalised to the average of the normoxia condition. Bars represent mean Log_2_(fold change) ± SD, n = 7 biological replicates. Statistical test: two-way ANOVA with Bonferroni’s multiple comparisons test. Gene expression was normalised to *ACTB* for RT-qPCR. Significance levels: *p < 0.05, **p < 0.01, ***p < 0.001.

**Figure S2.**
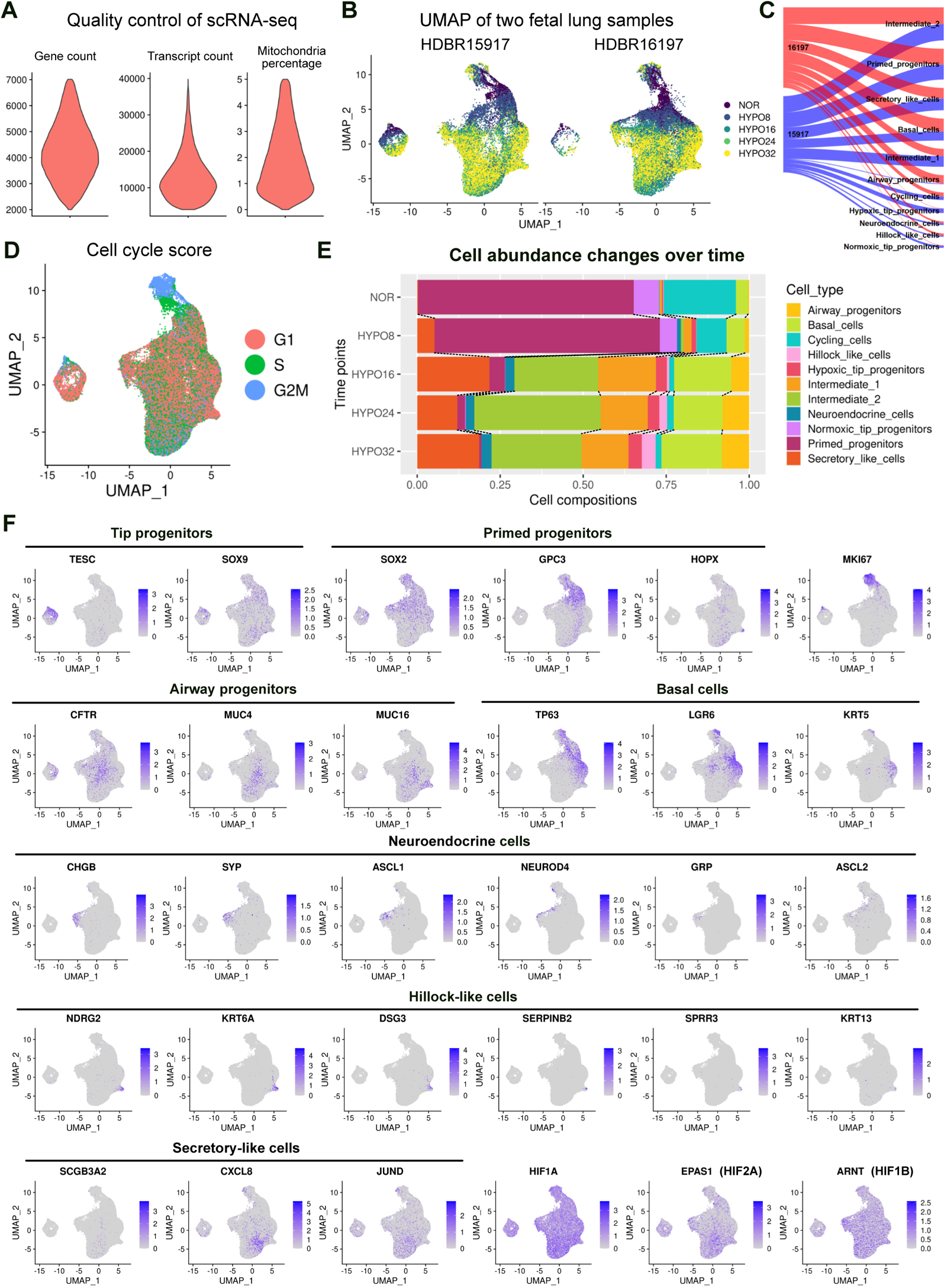
Characterisation of organoid single cell transcriptomic dataset. Related to Figure 2. (A) Quality control and cell filtering standards, showing the gene count, transcript count and mitochondrial gene percentage for filtered cells. (B) UMAP of cells sampled at different time points from two biological donors. (C) The contribution of two biological donors to the annotated cell types. (D) Cell cycle scores across all cells in the dataset. (E) Cell abundance changes of annotated cell types across different time points. (F) Feature plots showing marker gene expression patterns. The plot for each gene was scaled to maximum expression level of the gene.

**Figure S3.**
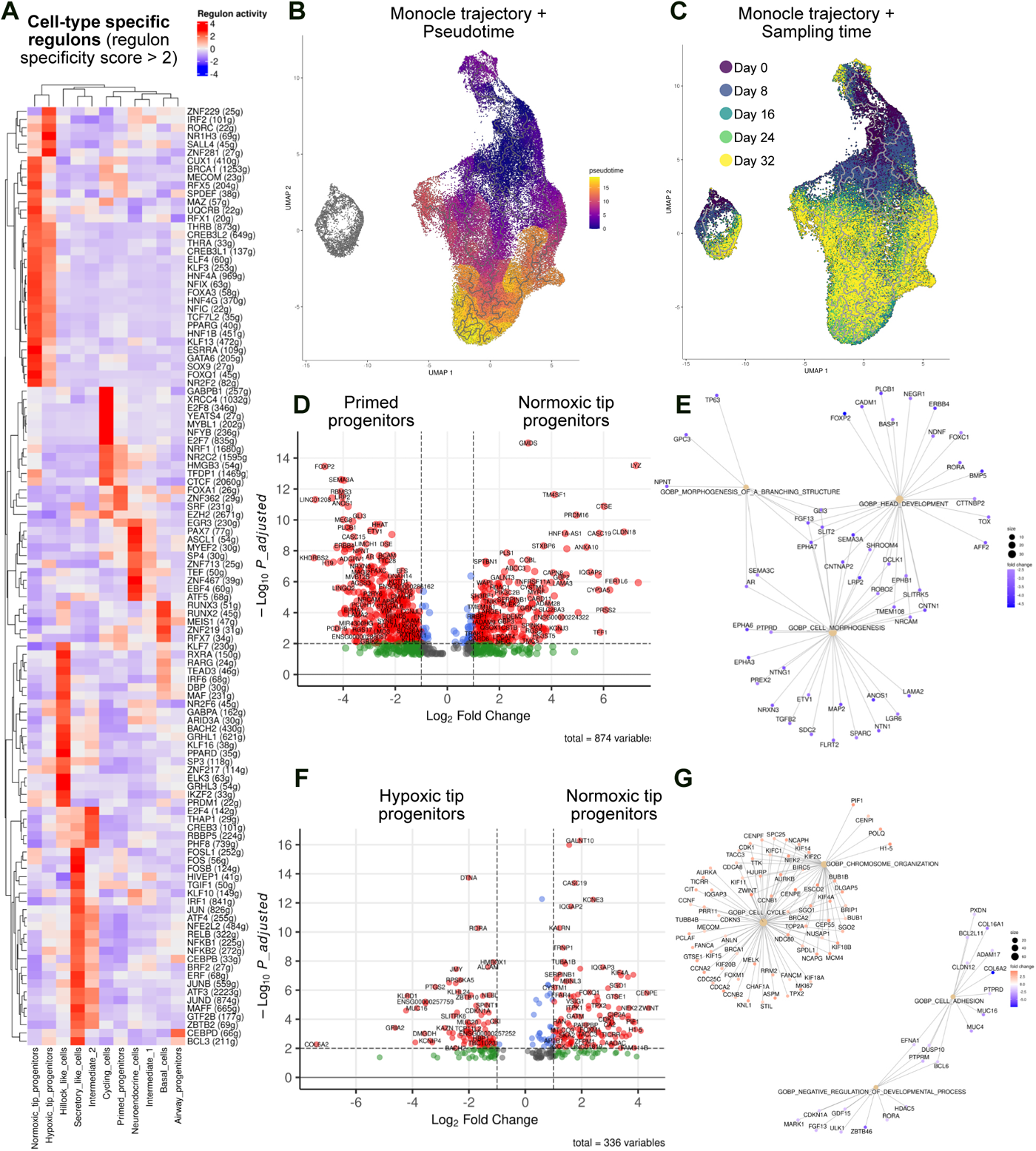
Regulons, trajectories, and differential gene expression analyses for organoid single cell transcriptomic dataset. Related to Figure 2. (A) Cell-type specific regulons (filtered by regulon specificity score > 2) identified with SCENIC. (B) and (C) Monocle 3 trajectories overlaying with pseudotime (B) or actual sampling time (C). (D) Volcano plot showing 874 filtered differentially expressed genes (*Padj* < 0.05) between primed progenitors and normoxic tip progenitors generated by pseudobulk analysis with MAST. (E) Linkage between DEGs in (D) and top-ranked Gene Ontology terms. (F) Volcano plot showing 336 filtered differentially expressed genes (*Padj* < 0.05) between hypoxic tip progenitors and normoxic tip progenitors generated by pseudobulk analysis with DESeq2. (G) Linkage between DEGs in (F) and top-ranked Gene Ontology terms.

**Figure S4.**
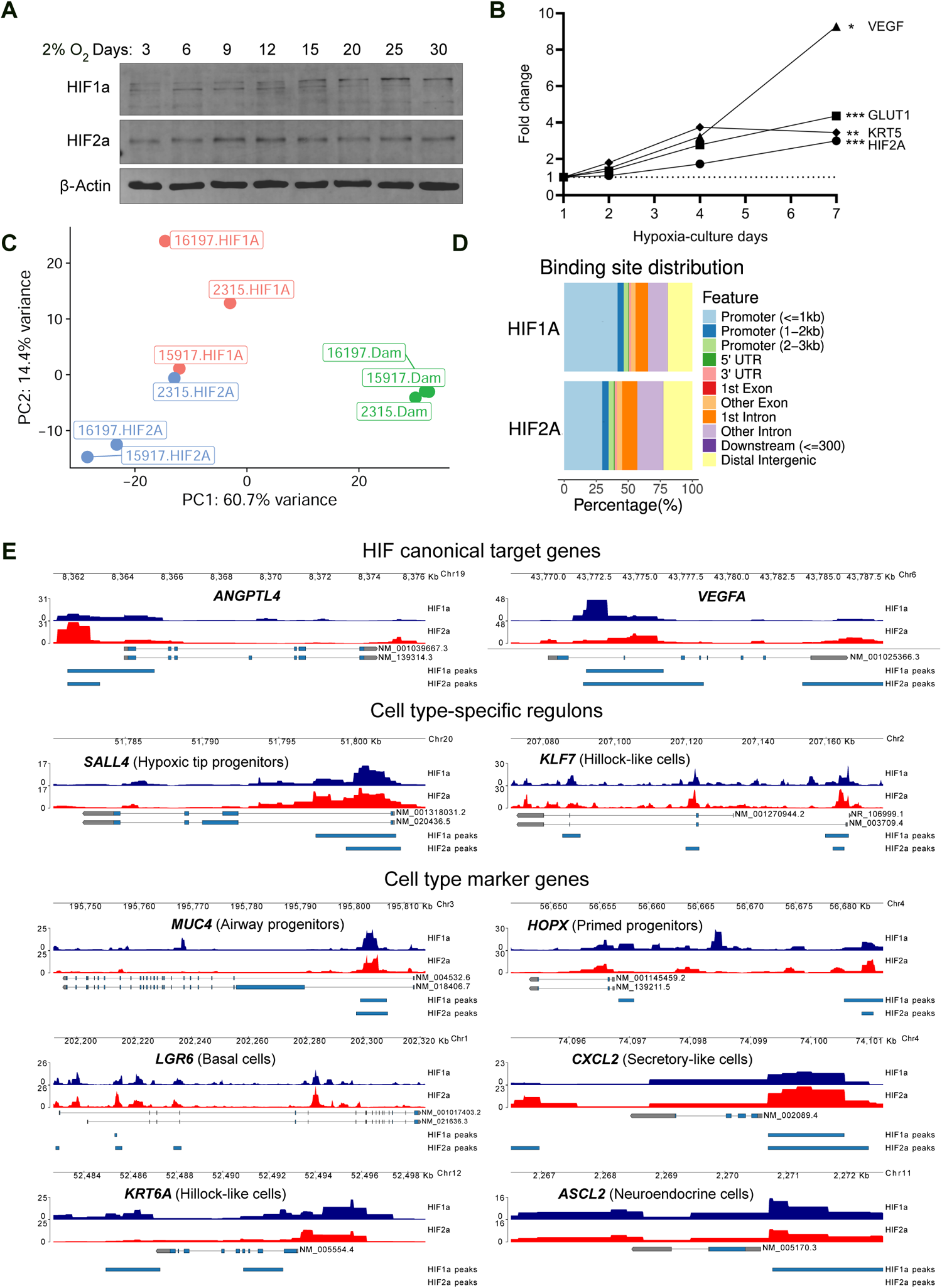
Activation of the HIF pathway and analyses of HIF1α and HIF2α targeted DamID-seq. Related to Figure 3. (A) Protein expression level of HIF1α, HIF2α and β-actin detected by Western blot in organoids cultured under hypoxia for 3-30 days. (B) RT-qPCR of organoids cultured under hypoxia for 1, 2, 4, 7 days. Fold changes were normalised to hypoxia day 1 expression levels. Data shown as mean fold change, n = 3 biological donors. The significance of the curve slopes differing from zero was tested by linear regression. Gene expression was normalised to *ACTB*. Significance levels: *p < 0.05, **p < 0.01, ***p < 0.001. (C) PCA plot of DamID samples from 3 organoid lines. (D) Categories of HIF1α and HIF2α binding sites. (E) Gene track views showing averaged DamID signals from three biological replicates over representative HIF1α and HIF2α target genes with consensus peaks labelled. The cell type markers and regulon transcription factors were selected from the organoid scRNA-seq dataset.

**Figure S5.**
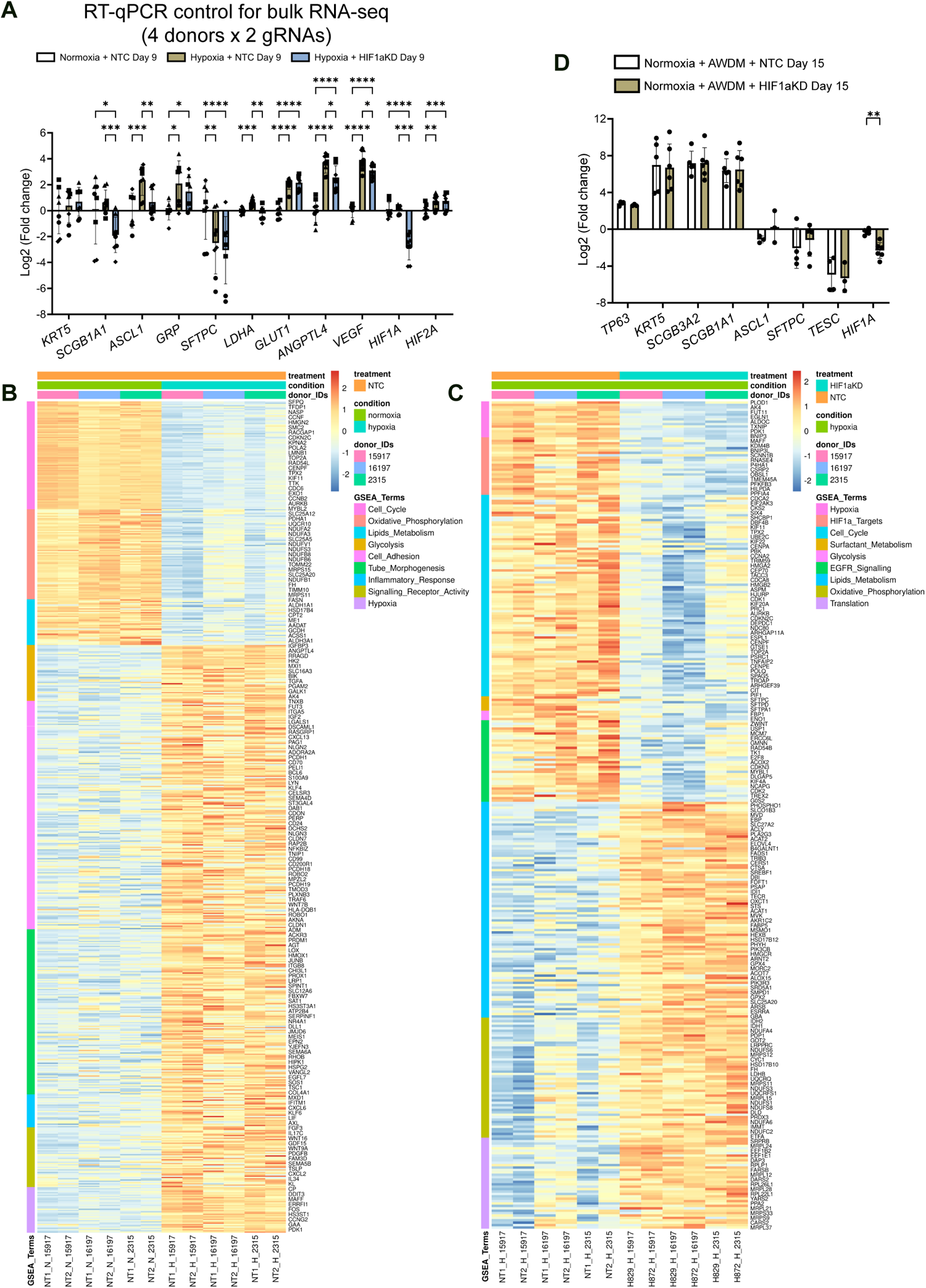
Bulk RNA-seq, differential gene expression analysis and airway differentiation tests for HIF1α-CRISPRi organoids. Related to Figure 4. (A) RT-qPCR control of samples used for bulk RNA-seq. NTC and *HIF1A*-knock down (*HIF1A-*KD) organoids were cultured under normoxia or hypoxia for 9 days. Fold changes were normalised to the mean of NTC + normoxia condition. Bars represent mean Log_2_(fold change) ± SD, n = 8 experimental replicates from 4 biological donors with 2 gRNAs. For bulk RNA-seq, 6 replicates (3 biological donors with 2 gRNAs) for each condition were selected. Statistical test: two-way ANOVA with Tukey’s multiple comparisons test. (B) Heatmap of 655 DEGs related to GSEA terms enriched in hypoxia compared to normoxia NTC organoids. Every 1 in 4 genes are labelled due to space limitations. (C) Heatmap of 345 DEGs related to GSEA terms enriched in *HIF1A-*KD compared to NTC hypoxic organoids. Every 1 in 2 genes were labelled due to space limitations. (D) RT-qPCR of NTC and *HIF1A-*KD organoids cultured in airway differentiation medium (AWDM) under normoxia for 15 days. The fold changes were normalised to the mean of normoxia + SRM condition (not shown). Bars represent Log_2_(fold change) ± SD, n = 5 (NTC), 6 (*HIF1A-*KD) experimental replicates from 3 biological donors with 2 gRNAs. Statistical test: two-way ANOVA with Bonferroni’s multiple comparisons test. Gene expression was normalised by *ACTB* in RT-qPCR. Significance levels: *p < 0.05, **p < 0.01, ***p < 0.001.

**Figure S6.**
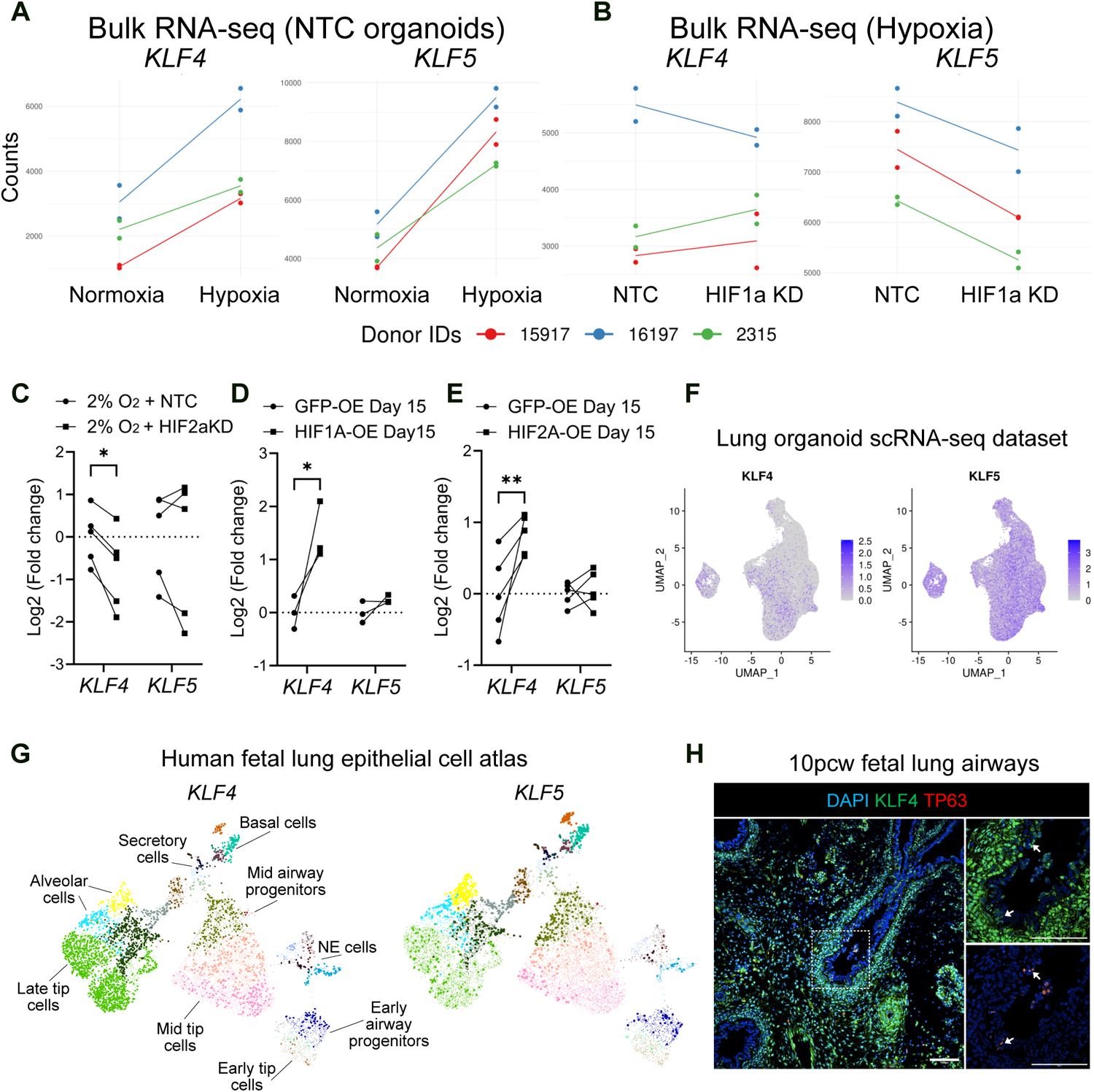
Expression regulations by hypoxia and expression patterns of KLF4 and KLF5. Related to Figure 6. (A) and (B) *KLF4* and *KL5* expression level from bulk RNA-seq data (as described in Figure 4) comparing Normoxia + NTC to Hypoxia + NTC (A), or Hypoxia + NTC to Hypoxia + *HIF1A*-knock down (KD) (B), n = 6 (2 gRNAs and 3 donors) for each condition. Each line indicates the change of the average counts of the two gRNA replicates from the same biological donor. (C) *HIF2A* knock down decreased *KLF4* but not *KLF5* expression under hypoxia. Data shown as Log_2_(fold change), n = 5 experimental replicates from 4 biological donors. 2 gRNAs used. (D) and (E) Stabilised HIF1α and HIF2α (as described in Figures 4,5) overexpression under normoxia increased *KLF4* expression. Data shown as Log_2_(fold change), n = 3 (HIF1α), 5 (HIF2α) biological donors. (F) *KLF4* and *KLF5* expression in organoid scRNA-seq dataset (as described in Figure 2). (G) *KLF4* and *KLF5* expression in epithelial cells of the human fetal lung atlas.^6^ (H) Immunostaining of 10 pcw human fetal lung section showing KLF4 and TP63 expression. Arrows indicate KLF4^+^TP63^+^ cells. Scale bars = 100 µm. RT-qPCR gene expression was normalised to *ACTB*. Statistical test: two-way ANOVA with Bonferroni’s multiple comparisons test. Significance levels: *p < 0.05, **p < 0.01.

